# CryoFlex characterizes structural motion between conformations directly from cryo-EM density maps

**DOI:** 10.64898/2026.09.01.748519

**Authors:** Hao Dong, Fuwei Li, Yuanbo Chen, Shuai Tang, Chenxuan Ji, Xinsheng Wang, Zhou Lu, Bin Hu, Fa Zhang, Xiaohua Wan

**Affiliations:** Key Laboratory of Brain Health Intelligent Evaluation and Intervention, Beijing Institute of Technology, Beijing, 100081, China; School of Medical Science and Engineering, Beijing Institute of Technology, Beijing, 100081, China; School of Artificial Intelligence, Beijing Institute of Technology, Beijing, 100081, China

**Keywords:** Cryo-EM, Computational Structural Biology, Non-rigid Registration

## Abstract

Heterogeneous cryo-EM reconstruction methods can resolve multiple conformational states, but understanding their functional implications often requires determining which regions move between states and quantifying their displacement magnitudes. Comparing atomic models can quantify motion, but flexible regions are often incompletely modeled. Direct map comparison is limited by variable map quality and ambiguous correspondence between displaced density features. Here, we present CryoFlex, a method that estimates the direction and magnitude of structural motion directly from reconstructed density maps of selected states. The resulting motion analysis supports functional interpretation and guides downstream structure processing. CryoFlex localized and quantified structural motion across diverse map pairs, including those with incomplete atomic-model coverage or weak local density. Against C*α* displacements, CryoFlex achieved an endpoint error of about 1 Å. In separate benchmarks, it outperformed registration baselines on clean and noisy inputs. Thus, CryoFlex complements heterogeneous reconstruction workflows with direct quantification of structural motion between reconstructed states.

## 1. Introduction

Heterogeneous reconstruction methods can resolve a range of conformational states from a single cryogenic electron microscopy (cryo-EM) dataset (Zhong et al., 2021; Punjani and Fleet, 2023). Conformational changes between these states are often closely linked to macromolecular function (Murata and Wolf, 2018). In the magnesium channel CorA, for example, a hinge-bending movement of the transmembrane helices opens the ion-conducting pore (Matthies et al., 2016). Reconstructing the states alone does not describe how structural regions move between two selected states. Analyzing these changes can identify the regions involved and describe how they move. Such analyses can help researchers interpret macromolecular function and inform other downstream tasks, for example, focused refinement in mobile regions (Nakane et al., 2018). However, characterizing structural motion between selected states remains challenging because macromolecular conformational changes can involve coupled motions across multiple regions and spatial scales (Punjani and Fleet, 2021, 2023).

A common way to characterize these changes is to build or fit an atomic model for each state (Brown et al., 2015; Afonine et al., 2018). The resulting models can then be compared through structural superposition, residue-level deviations or domain displacements (Taylor et al., 2014). This approach places conformational changes in a chemically interpretable coordinate system. Its reliability nevertheless depends on models that are sufficiently complete and accurate across both states. This requirement is hardest to satisfy in flexible regions, where conformational averaging lowers local resolution and blurs the density (Kucukelbir et al., 2014). Main-chain connectivity may consequently become ambiguous, leaving these regions incompletely modeled or unmodeled (Brown et al., 2015; Afonine et al., 2018). Direct comparison of density maps provides a complementary route, but noise and local-resolution variation can obscure density features. Visual overlays and difference maps can reveal where two reconstructions differ (Goddard et al., 2007; Pettersen et al., 2021). Local correlation analysis provides a quantitative measure of map agreement (Warshamanage et al., 2022). However, these analyses do not establish correspondences between displaced density features. They therefore cannot determine the direction or magnitude of the underlying structural displacement.

These limitations motivate direct comparison of reconstructed maps, particularly when atomic models are incomplete or unavailable. We consider two cryo-EM density maps that represent different conformations of the same macromolecular complex. These maps may be obtained with heterogeneous reconstruction methods (Zhong et al., 2021; Punjani and Fleet, 2023). Characterizing how structural regions move between the states requires a spatially coherent displacement field. Such a field establishes correspondences between density features and reports the direction and magnitude of local displacement. This analysis focuses on conformational differences between states.

Here, we present CryoFlex, a density-native approach for characterizing structural motion between selected states directly from cryo-EM density maps. CryoFlex estimates the direction and magnitude of structural motion without fitted atomic models. It also produces a voxel-aligned flexibility map that quantifies regional motion and provides spatial information for downstream structure processing. We first apply CryoFlex to NTCP, EMPIAR-10345, EMPIAR-10073 and CryoBench Spike, where it identifies gate opening and regional motions under incomplete atomic-model coverage or weak density. We then evaluate displacement accuracy and robustness on simulated TmrAB and PCAT1 maps across map resolutions and noise levels. Benchmark comparisons show improved displacement recovery, and ablation experiments confirm the contribution of the registration constraints.

## 2. Results

### 2.1. Overview of CryoFlex

CryoFlex estimates structural motion between a pair of aligned cryo-EM density maps representing two selected conformational states (Fig. 1). Density-aware sampling first converts each map into a density-aware point cloud that encodes local density information at the sampled points. CryoFlex then extracts keypoints and matches their features to establish sparse correspondences between the source and target point clouds.

**Figure 1.**
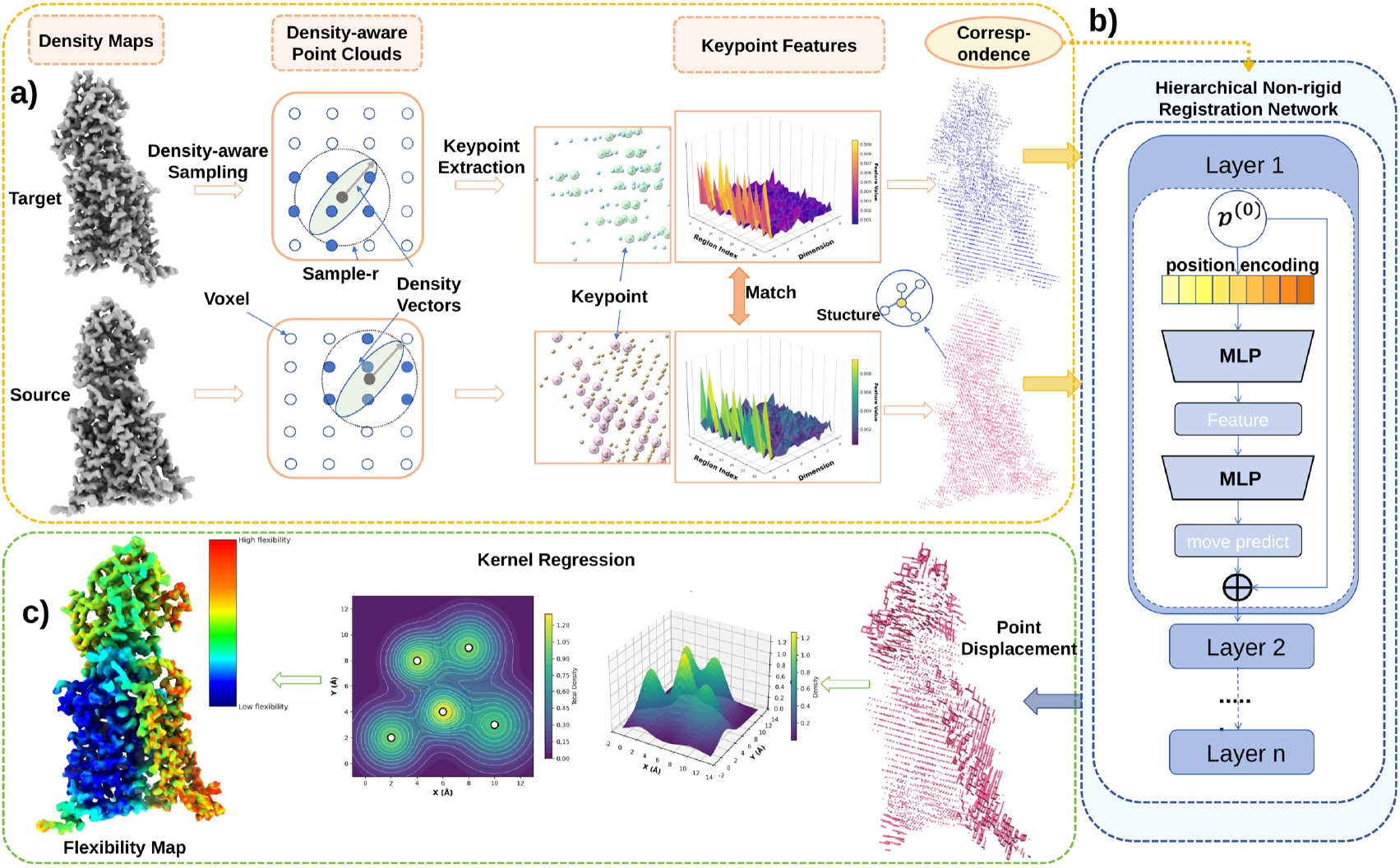
Overview of the CryoFlex workflow. **a**, CryoFlex takes aligned source and target cryo-EM density maps representing two selected conformational states. Density-aware sampling converts each map into a density-aware point cloud. Keypoints are extracted, and their features are matched to establish sparse correspondences between the source and target point clouds. **b**, A hierarchical non-rigid registration network estimates spatially coherent point displacements from the source point cloud toward the target. Sparse correspondences anchor the deformation, while local motion coherence regularizes neighboring point displacements. **c**, Kernel regression transfers the predicted point displacements to the source voxel grid, yielding a voxel-aligned displacement field whose magnitude defines the flexibility map.

A hierarchical non-rigid registration network estimates spatially coherent displacements that map the sampled source points toward the target point cloud. Sparse correspondences anchor the deformation, while local motion coherence regularizes neighboring point displacements. The resulting point displacements specify the direction and magnitude of the estimated motion.

Finally, kernel regression transfers the point displacements to the source voxel grid, yielding a voxel-aligned displacement field. The magnitude of this field defines the flexibility map, which reports the spatial distribution of between-state displacement magnitude. The displacement field and flexibility map support functional interpretation and downstream structure processing.

### 2.2. CryoFlex characterizes between-state motion in NTCP density maps

We first applied CryoFlex to two NTCP conformations (inward-facing, PDB 7PQG; open-pore, PDB 7PQQ) (Goutam et al., 2022). NTCP is a hepatocyte bile acid transporter and cellular entry receptor for hepatitis B and D viruses and undergoes a large pore-opening transition between the inward-facing and open-pore states. This density map pair provides a biologically interpretable case for mapping large-scale between-state motion. Overlays of the two density maps, shown in two orientations separated by a 45^◦^ rotation (Fig. 2a), highlight where the maps do not spatially coincide. However, these overlays do not assign correspondences between the two maps or provide a quantitative description of the local motion, including its direction, magnitude, or distribution across the transmembrane helices. From the same map pair, CryoFlex produces two outputs of a single density-native deformation: a displacement field on the source density map, illustrated by displacement vectors on a density-aware point cloud (Fig. 2b), and a flexibility map on the voxel grid that renders the corresponding displacement magnitude (Fig. 2c). Together, these outputs extend visual map comparison into a quantitative displacement description that localizes between-state displacement across the transmembrane helices without using atomic models to derive the deformation.

**Figure 2.**
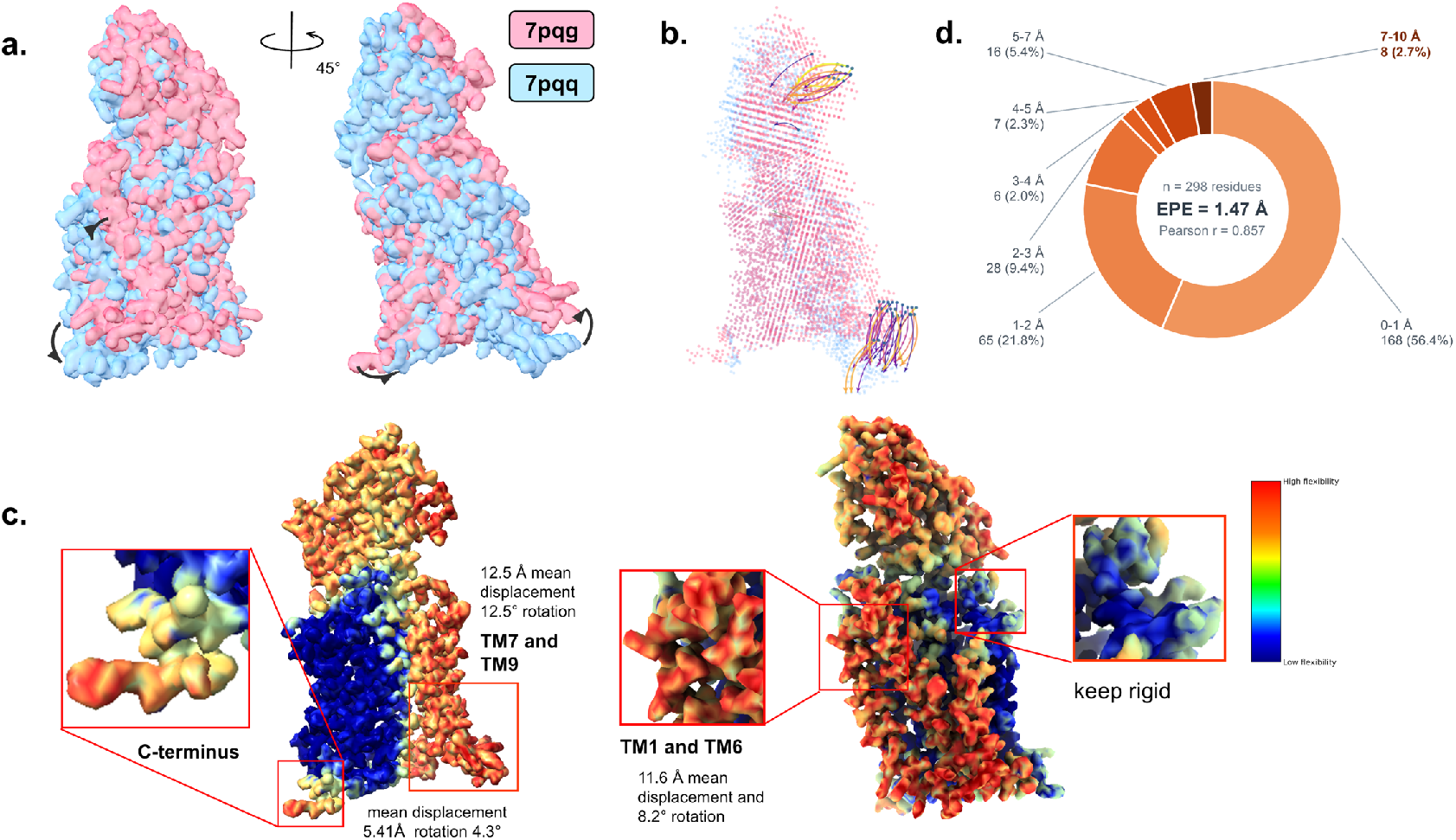
CryoFlex analysis of an NTCP density map pair. The inward-facing and open-pore states of the human sodium taurocholate cotransporting polypeptide (NTCP) are shown. **a**, Overlays of the inward-facing density map (pink; PDB 7PQG) and open-pore density map (blue; PDB 7PQQ), shown in two orientations separated by a 45^◦^ rotation. Black curved arrows schematically mark regions where the two overlaid maps visibly diverge and do not quantify motion. **b**, CryoFlex displacement vectors on the density-aware point cloud. Vectors are displayed in two representative mobile regions; other displacement vectors are omitted for clarity. **c**, Flexibility map on the voxel grid showing displacement magnitude, displayed in the same two orientations as in **a.** Insets report segment-level mean displacement and rotation: distal TM7/TM9, 12.5 Å and 12.5^◦^; TM1/TM6, 11.6 Å and 8.2^◦^; and the C-terminal segment, 5.4 Å and 4.3^◦^. The inset labeled “keep rigid” marks a representative low-displacement region. **d**, Consistency analysis for chain A (*n* = 298 residues). The donut chart shows the distribution of per-residue endpoint errors (EPE) between the CryoFlex displacement vectors sampled at C*α* positions and the C*α* displacement vectors between the paired atomic models (see Methods). Center: EPE = 1.47 Å and Pearson *r* = 0.857 between the corresponding displacement magnitudes; 87.6% of residues have an EPE below 3 Å, and 78.2% have an EPE below 2 Å.

We then analyzed the CryoFlex displacement field at the regional level, summarizing the dense motion information in terms of segment-level mean displacement and rotation. The central helical scaffold formed by TM2 to TM4 and the proximal parts of TM7 to TM9 is expected to maintain the transport pathway across states, whereas the panel domain helices TM1 and TM6 gate access to the extracellular entrance. The flexibility map is consistent with this expected gate-opening organization: it assigns low displacement magnitudes to this scaffold with a localized increase at the C-terminus of TM9, a region associated with pore dilation in the structural model, and highlights TM1 and TM6 as a mobile band along the extracellular entrance. Among the manually annotated helical segments, the distal parts of TM7 to TM9 showed the largest coordinated rearrangement (12.5 Å mean displacement, 12.5^◦^ rotation); TM1 and TM6 formed a second mobile band at the extracellular gate (11.6 Å, 8.2^◦^); and the C-terminal segment showed a smaller, localized displacement (5.4 Å, 4.3^◦^), whereas the central helices remained comparatively rigid (Fig. 2c, insets). Together, these motions are consistent with separation of the core and panel domains to open the transmembrane pathway for bile salt transport (Goutam et al., 2022). These regional summaries were computed from the CryoFlex displacement field and were not obtained by fitting the paired atomic models.

We next compared CryoFlex-derived local displacements with C*α* displacements measured from the paired atomic models. For chain A (*n* = 298 residues), we sampled the Gaussian-weighted CryoFlex displacement field at each C*α* position as defined in STAR Methods. Fig. 2d summarizes the distribution of per-residue endpoint errors (EPE) as a donut chart, with an EPE of 1.47 Å and a Pearson correlation of *r* = 0.857 between the corresponding displacement magnitudes reported at its center; in total, 87.6% of residues fall below 3 Å EPE and 78.2% below 2 Å. The largest residuals occur in the regions with the largest displacements, consistent with the attenuation expected when local motion is read from a voxel-averaged and kernel-regressed field. Fig. 2d provides an atomic-model consistency assessment of the CryoFlex displacement pattern across chain A.

Together, Fig. 2 shows how CryoFlex maps between-state motion in a density map pair through two density-native outputs: displacement vectors on the point cloud and a flexibility map on the voxel grid. Atomic-model C*α* displacement provided an external consistency reference, whereas CryoFlex derived the deformation exclusively from the two density maps. The following figures extend the same analysis to map pairs for which atomic-model references are incomplete or unavailable, where the density-native outputs can still be inspected directly on the maps.

### 2.3 CryoFlex characterizes motion where atomic-model coverage is incomplete

We next applied CryoFlex to two conformations of asymmetric *α*v*β*8 integrin reconstructed from the EMPIAR-10345 particle images in cryoSPARC (Campbell et al., 2020). The same dataset has also been used in prior continuous flexibility analysis (Punjani and Fleet, 2023), and we use it here to test density-native motion interpretation under incomplete model coverage. For this complex, available atomic models mainly describe the better resolved core region, whereas the flexible arm is poorly resolved or incompletely modeled in lower-resolution density (Cormier et al., 2018) (Fig. 3a). Prior work described this conformational change qualitatively but did not assign a displacement magnitude or rotation angle. EMPIAR-10345 provides a test case for density-native analysis under incomplete atomic-model coverage.

**Figure 3.**
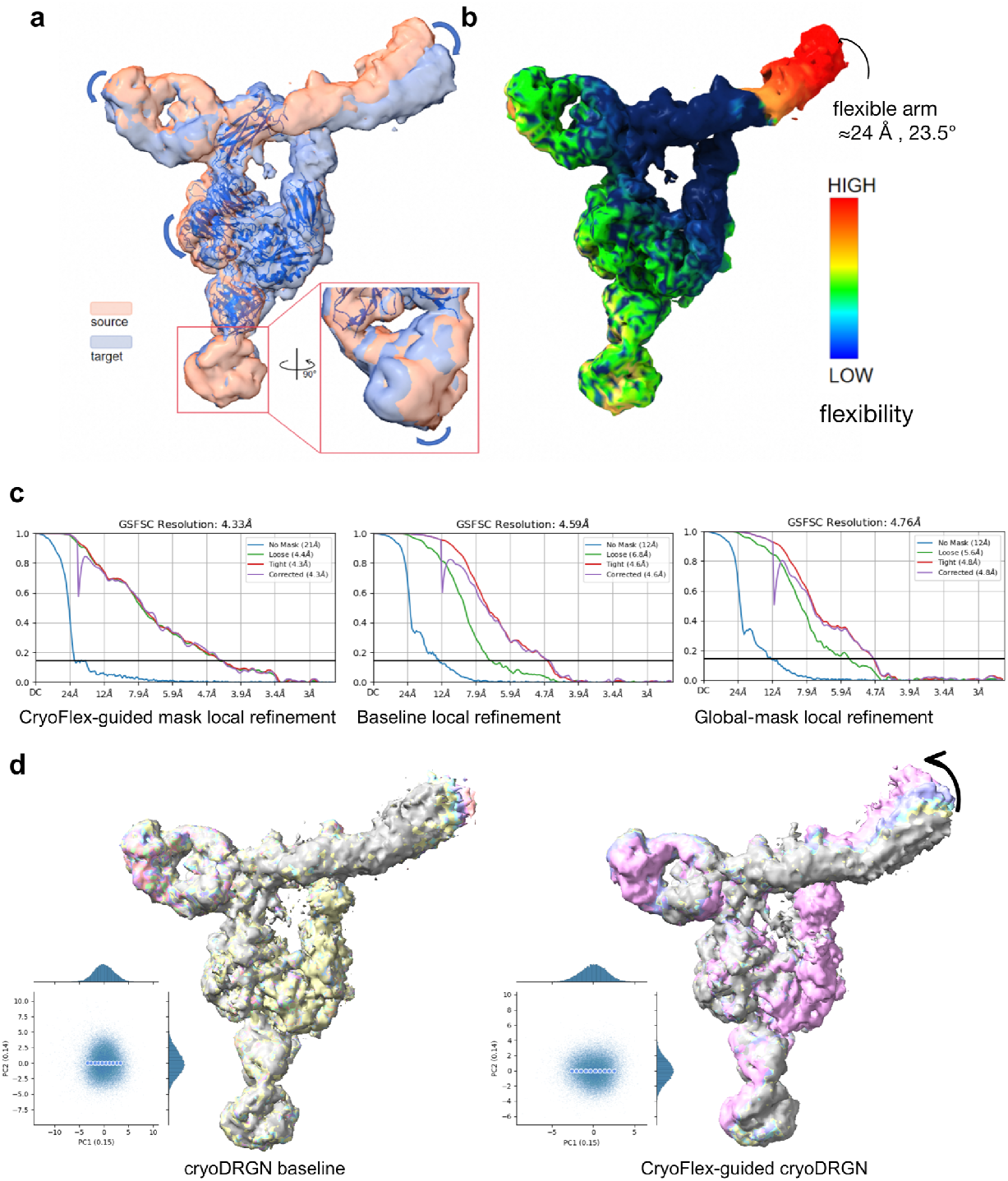
CryoFlex characterizes motion on EMPIAR-10345 (*α*v*β*8 integrin) when atomic-model coverage is incomplete. **a**, Overlay of the two *α*v*β*8 integrin conformations with the available atomic model, which mainly describes the better resolved core region while leaving the distal flexible arm incompletely modeled. **b**, Flexibility map on the voxel grid derived from the CryoFlex displacement field between the two density maps, with each voxel colored by displacement magnitude. The stable core and higher-displacement flexible arm are indicated. The arrow marks the dominant regional motion, and the inset reports approximately 24 Å mean displacement and 23.5^◦^ effective rotation relative to the core. **c**, Gold-standard FSC curves for three local refinement conditions using the same particle set and refinement settings: CryoFlex-guided (4.33 Å), original (4.59 Å) and global-mask (4.76 Å). **d**, Ten states sampled along PC1 and overlaid, comparing the cryoDRGN baseline (left) with cryoDRGN trained using spatially adaptive reference regularization guided by the CryoFlex flexibility map (right); each is shown with its PCA sampling inset.

CryoFlex generated a flexibility map on the voxel grid that separated the comparatively stable core region from a higher-displacement mobile region mapped to the flexible arm of the integrin (Fig. 3b). Each voxel in this map is colored by displacement magnitude, so the map reports the spatial distribution of between-state displacement magnitude rather than an intrinsic flexibility parameter. This spatial pattern is consistent with prior continuous flexibility analysis of the same dataset. We then summarized the CryoFlex displacement field within the mobile region by its mean displacement and rotation relative to the core region. The mobile region showed a mean displacement of approximately 24 Å and an effective rotation of 23.5^◦^, whereas the core region remained close to zero displacement (Fig. 3b, inset). The arrow in the panel denotes the dominant regional motion inferred from the displacement field, not an atom-by-atom trajectory. The flexible-arm motion is consistent with rotation of the headpiece about the *α*v knee, a conformational mode proposed to support ligand surveillance (Cormier et al., 2018). These results provide a quantitative, density-native characterization of the regional motion previously identified in this dataset, without requiring atomic models.

Because the flexibility map remains spatially aligned with the source density, we next tested its use as motion information in existing structureprocessing workflows. We first used the flexibility map to construct a focused mask for local refinement. With the particle set and refinement settings held constant, we compared the original, CryoFlex-guided and global-mask local refinement conditions (Fig. 3c). Without lowering the overall resolution, the CryoFlex-guided condition improved the local resolution of the mobile region, reaching the highest overall gold-standard FSC resolution of the three conditions at 4.33 Å, compared with 4.59 Å for the original condition and 4.76 Å for the global-mask condition. The automated mask-construction procedure is described in Sec. 8.5.

We next used the same flexibility map as a spatial prior during cryoDRGN training on the same particle set by applying the spatially adaptive reference regularization described in Sec. 8.5. Relative to the cryoDRGN baseline, the conformational series sampled along PC1 with this regularization showed better-defined core density, fewer fragmented density features and a more continuous progression in the flexible arm (Fig. 3d). Together, these analyses show that the voxel-aligned flexibility map can guide both focused refinement and heterogeneous reconstruction.

### 2.4. CryoFlex characterizes tri-snRNP head motion under reduced local structural definition

We next evaluated CryoFlex on the yeast U4/U6.U5 tri-snRNP dataset EMPIAR-10073, which has previously been used to analyze continuous structural flexibility (Nguyen et al., 2016; Punjani and Fleet, 2023). Reconstruction of the particle images in cryoSPARC yielded two major conformations at resolutions of 4.24 Å and 5.28 Å, which were used as the source and target states, respectively. Compared with the source map, the target map showed greater noise and substantially reduced local structural definition in the head region, with attenuated and directionally blurred density (Fig. 4a). These features make state-to-state deformation analysis challenging because structural displacement must be distinguished from changes in local density quality.

**Figure 4.**
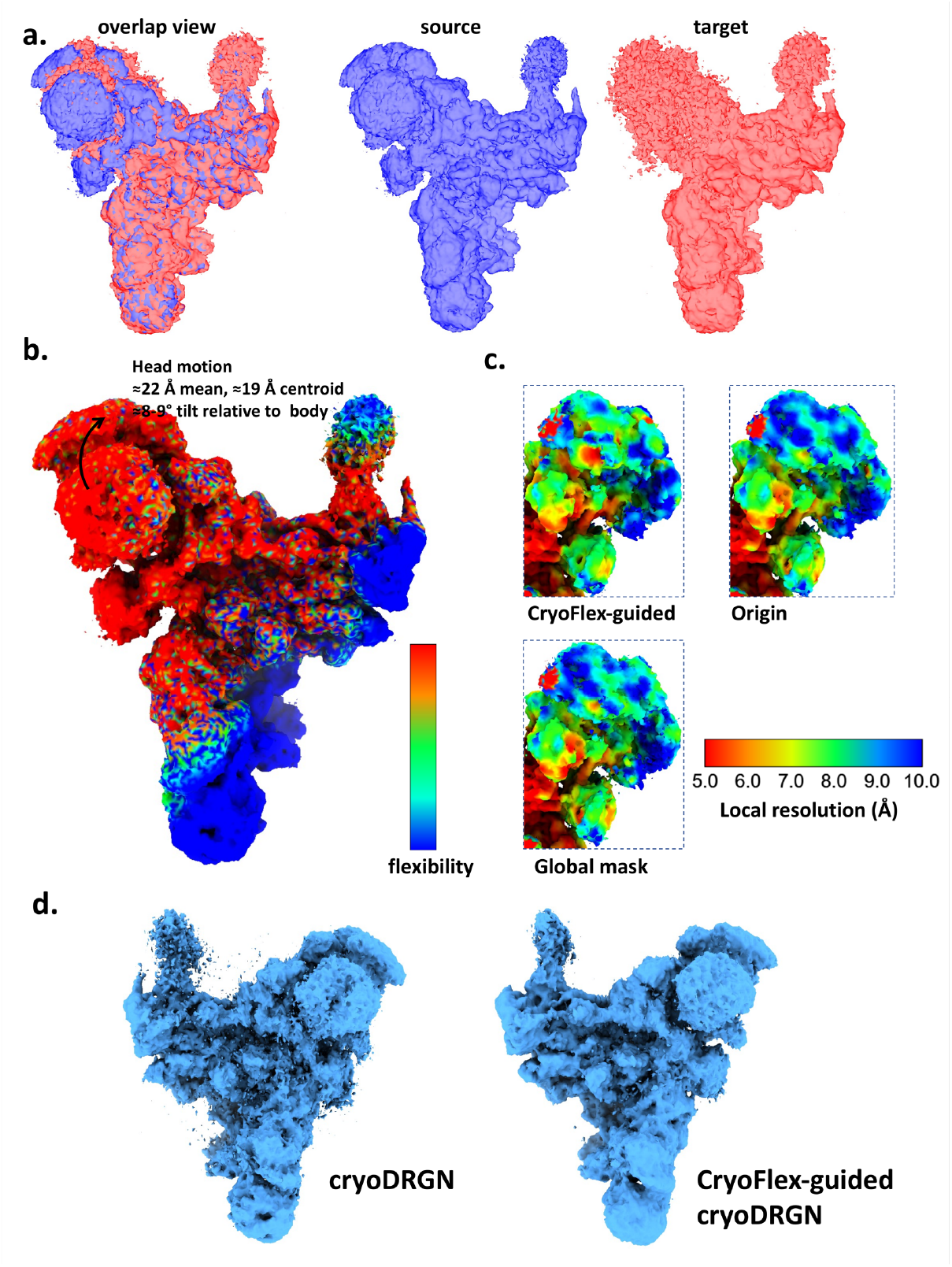
CryoFlex characterizes large-scale head motion in the yeast U4/U6.U5 tri-snRNP. **a**, Overlay and separate views of the 4.24 Å source map (blue) and 5.28 Å target map (red). Head density in the target map is attenuated and directionally blurred. **b**, CryoFlex flexibility map after alignment on the relatively stable body. The arrow summarizes the dominant head motion inferred from the displacement field. Regional analysis gives an approximately 22 Å mean displacement, 19 Å centroid displacement and 8–9^◦^ effective tilt of the head relative to the body. **c**, FSC-based local-resolution maps for the CryoFlex-guided, original and global-mask local refinement conditions, shown using the same color scale. All three conditions reached the same gold-standard FSC resolution of 4.24 Å, so the panels compare the spatial distribution of local resolution at matched overall resolution. **d**, Density maps obtained with baseline cryoDRGN (left) and cryoDRGN trained using the CryoFlex-guided spatial prior (right).

After alignment based on the relatively stable body region, CryoFlex identified a spatially coherent large-scale displacement of the head. We anatomically defined the head using Brr2 together with the Prp8 Jab1/MPN region in the public atomic model PDB 5GAN (Nguyen et al., 2016), rather than selecting the region solely from high values in the flexibility map. The head showed a mean displacement of approximately 22 Å and a centroid displacement of approximately 19 Å, together with an effective tilt of 8–9^◦^ relative to the body (Fig. 4b). This displacement estimate remained spatially coherent despite the attenuated and poorly defined density in the moving region.

The mobile head contains Brr2 and the Prp8 Jab1/MPN domain, placing the detected motion in a module involved in U4/U6 unwinding during spliceosome activation (Nguyen et al., 2016).

To assess the utility of this motion information in downstream structure-processing tasks, we applied the CryoFlex-derived flexibility map to local refinement and cryoDRGN reconstruction. With the particle inputs and refinement settings held constant, and without lowering the overall resolution (the original, CryoFlex-guided and global-mask conditions all reached the same gold-standard FSC resolution of 4.24 Å), FSC-based local-resolution maps showed higher estimated local resolution within the mobile region for the CryoFlex-guided condition (Fig. 4c). When incorporated as a spatial prior during cryoDRGN training, the CryoFlex flexibility map produced more continuous density, with fewer fragmented and scattered density features than the baseline reconstruction (Fig. 4d). Reduced local density quality did not prevent the flexibility map from guiding refinement and heterogeneous reconstruction.

### 2.5. CryoFlex characterizes Spike motion within a heterogeneous reconstruction workflow

We next applied CryoFlex to representative states generated by an existing heterogeneous reconstruction workflow. We used the CryoBench Spike dataset (Jeon et al., 2024) and selected three representative density maps from a cryoDRGN reconstruction (Zhong et al., 2021) that spanned progressive opening of the receptor-binding domain (RBD; Fig. 5a). These maps served as upstream inputs to CryoFlex and were analyzed as two adjacent pairs: map 1 to map 2 and map 2 to map 3. This sequential comparison allowed the conformational change to be examined across two adjacent intervals.

**Figure 5.**
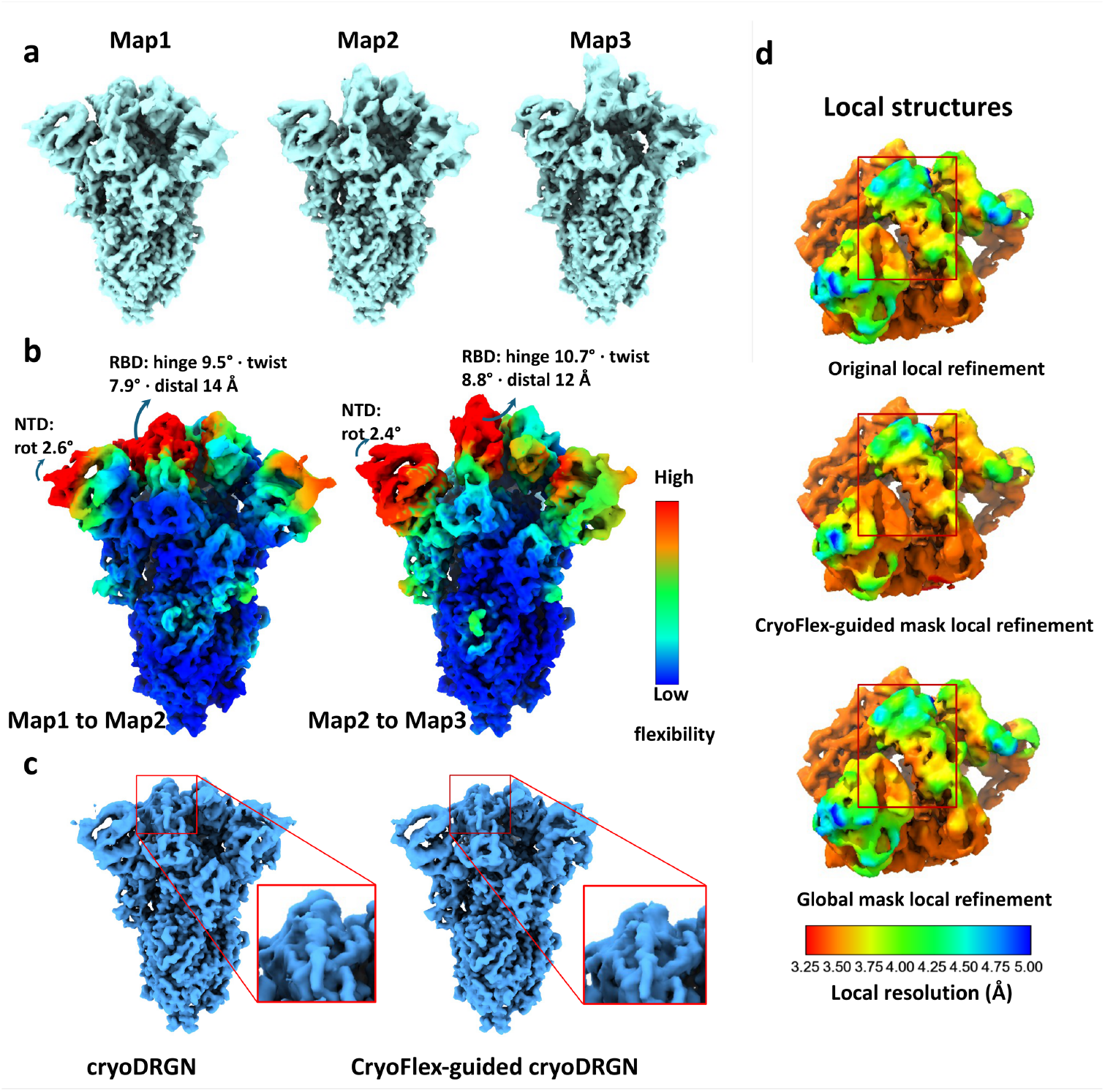
CryoFlex characterizes Spike motion within a heterogeneous reconstruction workflow. **a**, Three representative Spike density maps selected from a cryo-DRGN reconstruction, showing progressive opening of the receptor-binding domain (RBD) from map 1 to map 3. **b**, Flexibility maps on the voxel grid for the map 1 to map 2 (left) and map 2 to map 3 (right) transitions, colored by displacement magnitude from low (blue) to high (red). Arrows indicate the dominant regional motions. RBD labels report the hinge rotation, twist rotation and distal displacement: 9.5^◦^, 7.9^◦^ and 14 Å for map 1 to map 2; 10.7^◦^, 8.8^◦^ and 12 Å for map 2 to map 3. NTD labels report rotations of 2.6^◦^ and 2.4^◦^, respectively. **c**, Representative states sampled independently from latent spaces of baseline cryoDRGN and CryoFlex-guided cryoDRGN, trained with otherwise matched inputs and settings. Boxes mark the mobile region. **d**, Local structures after original, CryoFlex-guided and global-mask local refinement (top to bottom), colored by FSC-based local resolution. Red denotes lower Å values and higher resolution; blue denotes higher Å values and lower resolution. Corresponding gold-standard FSC curves are shown in Supplementary Fig. S3.

For the map 1 to map 2 transition, the flexibility map on the voxel grid localized the largest displacement to the opening RBD (Fig. 5b, left). This transition produced the larger distal displacement of the RBD and combined hinge and twist components in the displacement field. Regional analysis decomposed the RBD rotation into a hinge component of 9.5^◦^ and a twist component of 7.9^◦^. The mean displacement across the domain was 6.8 Å, while the displacement at the distal end reached approximately 14 Å. CryoFlex also identified spatially coherent upward displacement in two NTDs of the trimer. We analyzed the NTD carrying the larger motion (Fig. 5b), which rotated by 2.6^◦^, with a mean displacement of 2.1 Å and a distal displacement of approximately 4–5 Å. By comparison, the lower part of the trimer retained low displacement magnitudes, separating the mobile RBD and NTD from the more stable structural core.

The map 2 to map 3 comparison captured the adjacent interval of the opening progression (Fig. 5b, right). The RBD showed larger hinge and twist rotations than in the previous transition, whereas its distal displacement was smaller. The hinge and twist components were 10.7^◦^ and 8.8^◦^, respectively. The mean RBD displacement was 7.4 Å, and the displacement at its distal end was approximately 12 Å. The same NTD underwent a coupled but smaller motion, with a rotation of 2.4^◦^, a mean displacement of 2.8 Å, and a distal displacement of approximately 4–5 Å. The two pairwise analyses resolved distinct intervals of RBD opening: the first showed the larger distal displacement, whereas the second showed larger angular components. Together, these results are consistent with approximately rigid-body rotation of the RBD accompanied by smaller coordinated NTD motion.

The observed RBD opening moves the receptor-binding site toward an ACE2-accessible conformation required for receptor engagement (Wrapp et al., 2020).

We next incorporated the resulting spatial motion information into existing reconstruction and refinement workflows. CryoDRGN training was augmented with the spatially adaptive reference regularization described in Sec. 8.5, with the inputs, training parameters and processing settings otherwise held constant. Representative states were selected independently from the baseline and regularized latent spaces for comparison. The two reconstructions showed comparable overall density quality, while the reconstruction obtained with this regularization displayed more continuous density in the mobile region (Fig. 5c). We then used the CryoFlex-defined mobile region to guide local refinement and compared the resulting structures across the original, CryoFlex-guided and global-mask local refinement conditions. Without lowering the overall resolution (3.06 Å for all three conditions; Supplementary Fig. S3), FSC-based local-resolution maps showed higher estimated local resolution within the mobile region for the CryoFlex-guided condition (Fig. 5d). CryoFlex can analyze maps from heterogeneous reconstruction and return spatial motion information to reconstruction and refinement workflows.

To assess whether reconstruction-specific variation could be misinterpreted as conformational motion, we also analyzed independently reconstructed same-state half-map pairs for EMPIAR-10345 and Spike. These controls produced negligible or low-amplitude, spatially isolated displacement signals, in contrast to the stronger and spatially coherent patterns observed for the corresponding conformational-transition pairs (Supplementary Fig. S1).

### 2.6. CryoFlex estimates agree with C*α*-referenced displacements across nominal resolutions

This experiment tested whether the displacement vectors and magnitudes estimated by CryoFlex agreed with the C*α*-referenced endpoint displacement between two states, and whether that agreement was maintained as the nominal resolution of the input maps varied. Complete per-atom displacement references are generally unavailable for independently reconstructed experimental maps, because atomic-model coverage, local model confidence and residue correspondence can differ between states. We simulated source and target density maps independently from the pre-aligned paired atomic structures of TmrAB (6RAH → 6RAI) and PCAT1 (7T55 → 7T57) with `e2pdb2mrc` in EMAN2 (Tang et al., 2007) at a voxel size of 1.5 Å, each at nominal resolutions of 3, 4, 5, 6 and 8 Å. Because every map was generated from its own atomic state, the reference was the endpoint displacement vector between corresponding C*α* atoms. For each condition, the CryoFlex displacement vector was sampled at every source-state C*α* position. Vector errors and magnitude agreement were then evaluated against the corresponding C*α* endpoint displacement, as defined in Sec. 8.6. All 1,277 TmrAB and 1,464 PCAT1 C*α* atoms were retained for evaluation.

CryoFlex closely tracked the C*α* displacement references in both benchmark systems across all tested nominal resolutions (Fig. 6a,b). Agreement was quantified at two levels: the endpoint error and the RMSE, both computed between the estimated and reference displacement vectors, and the Pearson correlation coefficient, computed between their magnitudes. For TmrAB, the EPE ranged from 0.972 to 1.122 Å, the RMSE from 1.112 to 1.312 Å, and the Pearson correlation coefficient from 0.956 to 0.973. For PCAT1, the corresponding ranges were 0.954–1.068 Å, 1.139–1.265 Å and 0.974–0.979, respectively. Across two systems with distinct conformational changes, CryoFlex achieved an EPE of approximately 1 Å while tracking C*α* displacement magnitudes extending from sub-Å changes to displacements greater than 10 Å, with Pearson correlations above 0.95.

**Figure 6.**
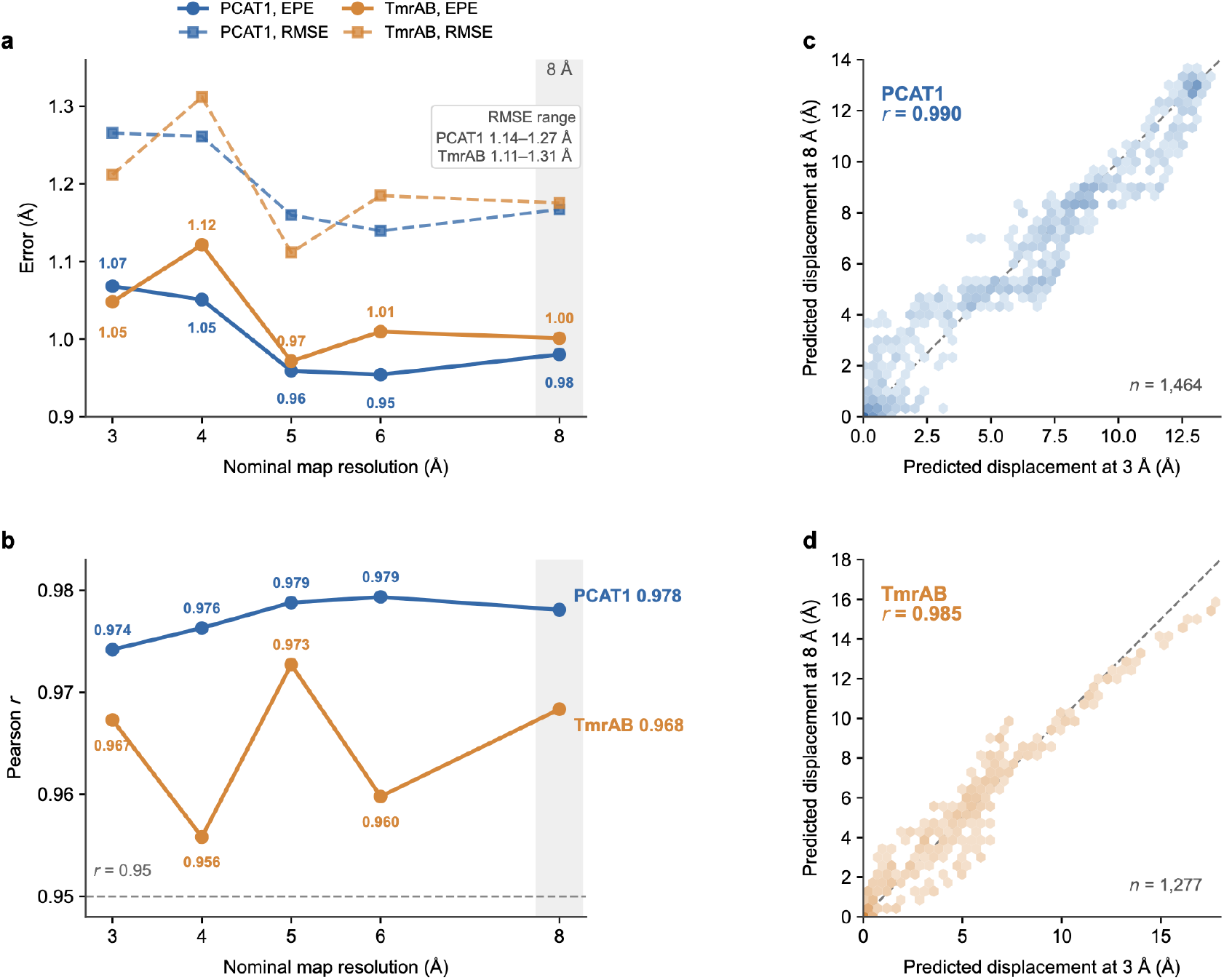
C*α*-referenced displacement accuracy and robustness across nominal resolutions. **a**, Endpoint error (EPE; solid lines and circles) and root mean squared error (RMSE; dashed lines and squares) between the CryoFlex-derived and corresponding C*α* displacement vectors for TmrAB and PCAT1. Source and target density maps were simulated independently from the paired atomic structures at nominal resolutions of 3, 4, 5, 6 and 8 Å. Numbers denote EPE values, the inset gives the RMSE range, and the shaded band marks the 8 Å condition. **b**, Pearson correlation between predicted and reference C*α* displacement magnitudes at each nominal resolution. The dashed horizontal line marks *r* = 0.95. **c,d**, Predicted displacement magnitudes at nominal resolutions of 3 Å (*x* axis) and 8 Å (*y* axis) for PCAT1 (**c**) and TmrAB (**d**). Hexagonal bins group C*α* positions, with darker colors indicating more atoms; the dashed diagonal denotes equality. Pearson correlations report cross-resolution stability rather than accuracy against the atomic-model reference. *n* = 1,277 matched C*α* atoms for TmrAB and *n* = 1,464 for PCAT1.

This C*α*-referenced accuracy did not deteriorate systematically as the nominal resolution varied from 3 to 8 Å. At a nominal resolution of 8 Å, CryoFlex retained EPEs of 1.001 Å for TmrAB and 0.980 Å for PCAT1, with Pearson correlations of 0.968 and 0.978, respectively. Moreover, the per-C*α* displacement magnitudes predicted at nominal resolutions of 3 and 8 Å were highly concordant, with cross-resolution correlations of 0.985 for TmrAB and 0.990 for PCAT1 (Fig. 6c,d). These results show that both the atomic-reference accuracy and the relative spatial pattern of the predicted displacement were maintained across the tested nominal-resolution range.

CryoFlex modestly underestimated the C*α* displacement magnitudes, with signed biases of the estimated magnitudes of approximately −0.81 to −0.98 Å for TmrAB and −0.52 to −0.58 Å for PCAT1. This systemdependent underestimation did not increase systematically as the nominal resolution became coarser. Together, these results show that CryoFlex captures the broad displacement range of both datasets with an EPE of approximately 1 Å against the C*α*-referenced endpoint displacement, and that this performance remained stable across nominal resolutions of 3–8 Å. The present analysis does not directly evaluate mismatched source–target resolutions, spatially varying local resolution or resolution anisotropy.

### 2.7. CryoFlex improves displacement recovery at sampled points and on the voxel grid

The aim of these benchmarks was to compare CryoFlex with representative registration methods against a simulation-defined voxel displacement field, and to assess how input noise affected the magnitude accuracy and spatial localization of CryoFlex predictions as well as their performance relative to the other methods. Two benchmarks were derived from TmrAB (6RAH → 6RAI) and PCAT1 (7T55 → 7T57), which represent a gate-like closure and a multi-domain rotation, respectively. In each benchmark, the displacement field relating the two maps was defined by simulation on the voxel grid, and its values at the source sampled points provided the reference displacement vectors and correspondence labels. These labels were used only for evaluation; all methods performed registration without access to them.

We evaluated two error measures at the sampled points: RMSE, which uses the known correspondence labels to measure whether each sampled point, and the corresponding local structural region, is displaced toward its correct target location; and RMSENN, which measures the nearest-neighbor distance between the deformed source point set and the target point set without using correspondence labels, reflecting predicted-to-target geometric proximity after registration. We refer to this correspondence-based displacement RMSE at the sampled points as point RMSE, and to the RMSE between the predicted and ground-truth displaced positions of each voxel (introduced below) as map RMSE. We compared against nine representative baselines spanning classical and learning-based non-rigid registration: CPD (Myronenko and Song, 2010), BCPD (Hirose, 2020), ICP (Rusinkiewicz and Levoy, 2001), NICP (Amberg et al., 2007), FilterReg (Gao and Tedrake, 2019), PSR (Zhao et al., 2024), Sinkhorn (Feydy et al., 2019), Nerfies (Park et al., 2021), and NDP (Li and Harada, 2022).

Point RMSE directly evaluates the displacement vectors at sampled points that form the basis of the flexibility map. CryoFlex achieved the lowest point RMSE and RMSENN on both benchmarks, with NDP as the closest baseline in this comparison (Table 1). Compared with NDP, CryoFlex reduced point RMSE from 0.83 to 0.59 on TmrAB and from 0.90 to 0.56 on PCAT1. RMSENN was also reduced from 0.68 to 0.46 and from 0.83 to 0.50, respectively.

**Table 1.** Point cloud registration accuracy on the simulation-defined benchmarks. Point RMSE is the correspondence-based error against the ground-truth displacement vectors at sampled points, computed using known correspondence labels. RMSENN is the nearest-neighbor geometric error between the deformed source point set and the target point set, which does not use correspondence labels. Both are evaluated on the two simulation-defined benchmarks. All methods received the same source and target point sets. Best value per column in bold; lower is better. Point cloud distance errors are reported in Å; the voxel grid is used as the common coordinate domain.

| Method | TmrAB (6RAH → 6RAI) | | PCAT1 (7T55 → 7T57) | |
| --- | --- | --- | --- | --- |
| | Point RMSE ↓ | RMSENN ↓ | Point RMSE ↓ | RMSENN ↓ |
| CPD | 3.54 | 1.71 | 4.71 | 2.40 |
| BCPD | 6.07 | 2.77 | 6.06 | 2.78 |
| ICP | 3.46 | 1.76 | 3.71 | 1.96 |
| NICP | 2.56 | 0.89 | 4.13 | 1.36 |
| FilterReg | 3.83 | 2.12 | 5.88 | 2.65 |
| PSR | 2.03 | 0.92 | 2.85 | 1.18 |
| Sinkhorn | 1.90 | 1.09 | 2.45 | 1.66 |
| Nerfies | 2.22 | 1.19 | 1.21 | 0.98 |
| NDP | 0.83 | 0.68 | 0.90 | 0.83 |
| **CryoFlex** | **0.59** | **0.46** | **0.56** | **0.50** |

We then asked whether the more accurate displacement vectors at sampled points were preserved after projection to the source-map voxel grid. For each method, these displacement vectors were projected onto the voxel grid using the same kernel regression procedure, giving the displacement field whose magnitude is the flexibility map, and were evaluated against the corresponding ground-truth displacement field (Table 2). CryoFlex achieved the lowest map RMSE and RMSE@15 and the highest Pearson correlation and Dice@15 on both benchmarks. Relative to NDP, CryoFlex reduced map RMSE from 0.51 to 0.37 on TmrAB and from 0.52 to 0.38 on PCAT1. It also gave the highest overlap with the ground-truth mask of the top 15% highest-displacement voxels, with Dice@15 values of 0.959 and 0.962 on TmrAB and PCAT1, respectively. These results show that the improvement in point-level displacement accuracy was retained after voxel-grid projection and was reflected in more accurate localization of high-displacement voxels. Complete point registration and flexibility map metrics for all evaluated methods are reported in Supplementary Tables S1 and S2.

**Table 2.** Voxel-grid displacement accuracy against the simulation-defined ground truth. Each method’s displacement vectors at sampled points are passed through the same kernel regression to form a displacement field on the voxel grid, whose magnitude is the flexibility map. Map RMSE measures the distance between the predicted and ground-truth displaced positions of each voxel. Pearson correlation (Corr) compares the predicted and ground-truth displacement magnitudes. Two further metrics are restricted to the ground-truth mask of the top 15% highest-displacement voxels: mask overlap (Dice@15) and displacement error within that mask (RMSE@15). Best value per column in bold. Map RMSE and RMSE@15 are in Å; Corr and Dice@15 are dimensionless. Baselines and inputs match Table 1.

| Method | TmrAB (6RAH → 6RAI) | | | | PCAT1 (7T55 → 7T57) | | | |
| --- | --- | --- | --- | --- | --- | --- | --- | --- |
| | Map RMSE ↓ | Corr ↑ | Dice@15 ↑ | RMSE@15 ↓ | Map RMSE ↓ | Corr ↑ | Dice@15 ↑ | RMSE@15 ↓ |
| CPD | 3.45 | −0.618 | 0.000 | 8.63 | 4.70 | −0.400 | 0.041 | 8.88 |
| BCPD | 3.32 | 0.708 | 0.695 | 2.95 | 3.15 | 0.408 | 0.240 | 2.00 |
| ICP | 3.07 | 0.444 | 0.461 | 7.68 | 2.28 | 0.547 | 0.388 | 5.26 |
| NICP | 1.94 | 0.852 | 0.672 | 4.89 | 3.04 | 0.363 | 0.369 | 6.30 |
| FilterReg | 2.94 | 0.151 | 0.292 | 7.16 | 3.37 | 0.381 | 0.313 | 2.68 |
| PSR | 0.92 | 0.953 | 0.886 | 1.52 | 1.42 | 0.835 | 0.782 | 3.05 |
| Sinkhorn | 0.87 | 0.972 | 0.905 | 1.96 | 1.37 | 0.876 | 0.893 | 2.54 |
| Nerfies | 1.67 | 0.930 | 0.766 | 4.08 | 0.91 | 0.930 | 0.901 | 1.65 |
| NDP | 0.51 | 0.986 | 0.946 | 0.87 | 0.52 | 0.975 | 0.960 | 0.77 |
| **CryoFlex** | **0.37** | **0.992** | **0.959** | **0.74** | **0.38** | **0.986** | **0.962** | **0.52** |

We next examined how reduced map quality affected the magnitude accuracy and spatial localization of CryoFlex predictions, and whether their relative advantage over the other methods was retained. Noise was added to both the source and target density maps before point sampling, thereby perturbing the source and target point sets sampled from the maps while preserving a common procedure for input generation and evaluation across methods. This perturbation design tested robustness to noise propagated from the density maps into the sampled point sets. We repeated the evaluation of the flexibility maps on the voxel grid at SNRs of 5, 2 and 1, with lower SNR indicating stronger noise (Table 3). To keep the comparison in the main text focused, we report CryoFlex together with two representative baselines: NDP, which was the closest baseline in the comparison on clean inputs, and Sinkhorn, which was consistently competitive under noisy conditions.

**Table 3.**
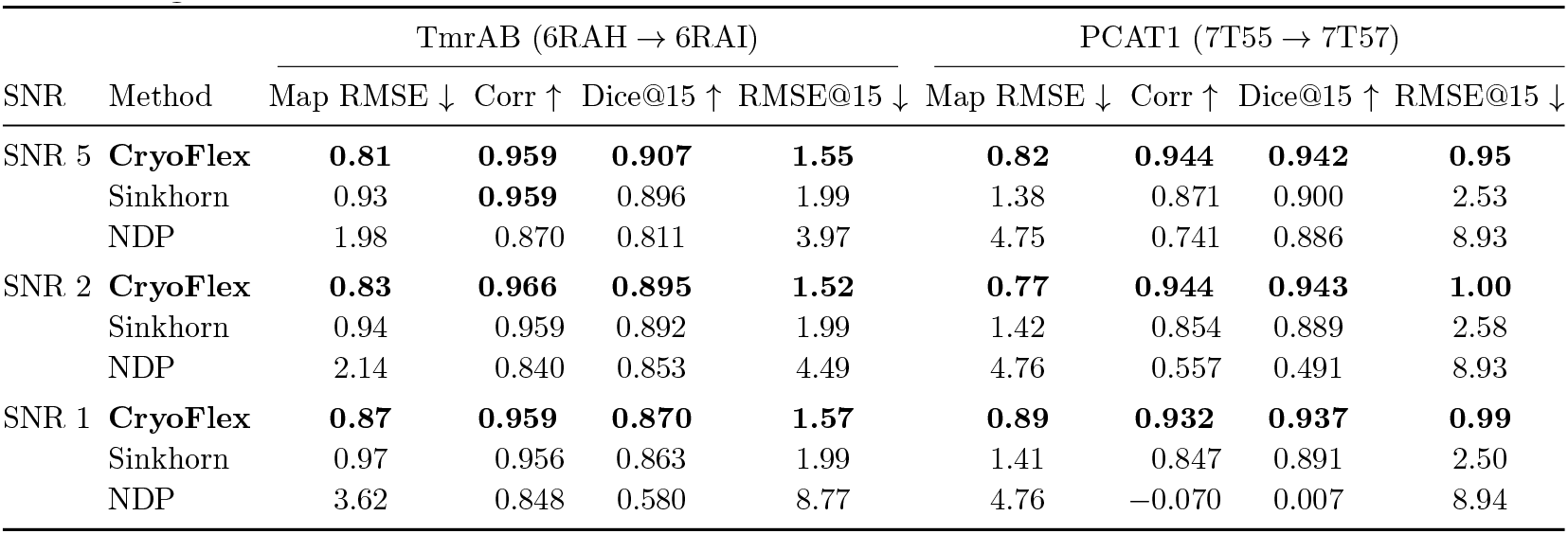
Voxel-grid displacement accuracy under noisy inputs. The evaluation in Table 2 is repeated at three signal-to-noise ratios (SNR 5, 2 and 1), where a lower SNR denotes stronger noise in the map inputs. Metrics and ground truth match Table 2: map RMSE and RMSE@15 are in Å, Corr and Dice@15 are dimensionless. For readability, CryoFlex is compared against two representative baselines: NDP, which was the closest baseline in the comparison on clean inputs, and Sinkhorn, which was consistently competitive under noisy conditions; the full method list is reported in the Supplementary. Best value per column within each SNR is shown in bold; lower map RMSE and RMSE@15 are better, higher Corr and Dice@15 are better.

| SNR | Method | TmrAB (6RAH → 6RAI) | | | | PCAT1 (7T55 → 7T57) | | | |
| --- | --- | --- | --- | --- | --- | --- | --- | --- | --- |
| | | Map RMSE ↓ | Corr ↑ | Dice@15 ↑ | RMSE@15 ↓ | Map RMSE ↓ | Corr ↑ | Dice@15 ↑ | RMSE@15 ↓ |
| SNR 5 | **CryoFlex** | **0.81** | **0.959** | **0.907** | **1.55** | **0.82** | **0.944** | **0.942** | **0.95** |
|  | Sinkhorn | 0.93 | **0.959** | 0.896 | 1.99 | 1.38 | 0.871 | 0.900 | 2.53 |
|  | NDP | 1.98 | 0.870 | 0.811 | 3.97 | 4.75 | 0.741 | 0.886 | 8.93 |
| SNR 2 | **CryoFlex** | **0.83** | **0.966** | **0.895** | **1.52** | **0.77** | **0.944** | **0.943** | **1.00** |
|  | Sinkhorn | 0.94 | 0.959 | 0.892 | 1.99 | 1.42 | 0.854 | 0.889 | 2.58 |
|  | NDP | 2.14 | 0.840 | 0.853 | 4.49 | 4.76 | 0.557 | 0.491 | 8.93 |
| SNR 1 | **CryoFlex** | **0.87** | **0.959** | **0.870** | **1.57** | **0.89** | **0.932** | **0.937** | **0.99** |
|  | Sinkhorn | 0.97 | 0.956 | 0.863 | 1.99 | 1.41 | 0.847 | 0.891 | 2.50 |
|  | NDP | 3.62 | 0.848 | 0.580 | 8.77 | 4.76 | -0.070 | 0.007 | 8.94 |

The principal effect of noise on CryoFlex was an increase in displacement magnitude error, whereas the spatial localization of regions with large displacement was better preserved. The map RMSE increased from 0.37 to 0.81–0.87 on TmrAB and from 0.38 to 0.77–0.89 on PCAT1, and the RMSE@15 over the ground-truth top 15% voxels with the largest displacement increased from 0.74 to 1.52–1.57 and from 0.52 to 0.95–1.00, respectively. By comparison, Dice@15 decreased from 0.959 to 0.870–0.907 on TmrAB and from 0.962 to 0.937–0.943 on PCAT1, and the Pearson correlation remained high (0.959–0.966 and 0.932–0.944). Despite the decrease in absolute accuracy, CryoFlex achieved the lowest map RMSE and RMSE@15 and the highest Dice@15 on both benchmarks at all three SNR levels. At the most challenging setting, SNR 1, CryoFlex reduced the map RMSE relative to Sinkhorn from 0.97 to 0.87 on TmrAB and from 1.41 to 0.89 on PCAT1, while increasing Dice@15 from 0.863 to 0.870 and from 0.891 to 0.937, respectively. Noise increased absolute displacement errors, whereas the spatial localization of high-displacement regions was better preserved. CryoFlex also retained its relative advantage across the tested SNR range. Several baselines showed substantially larger degradation under these conditions, with complete results reported in Supplementary Tables S3–S5.

Together, these controlled evaluations provide quantitative evidence that CryoFlex improves both displacement recovery at sampled points and the accuracy of flexibility maps on the two simulation-defined benchmarks, which represent distinct large-scale motion patterns, and that its relative advantage was maintained across the tested SNR range.

### 2.8. CryoFlex registration constraints make complementary contributions to displacement accuracy

We next examined whether the performance advantage observed on the simulation-defined benchmarks depended on the CryoFlex-specific constraints. Using the clean TmrAB (6RAH → 6RAI) and PCAT1 (7T55 → 7T57) datasets, we kept the hierarchical deformation backbone, sampled point sets, network initialization procedure, optimization settings and point-to-voxel kernel regression fixed across variants. We compared the full CryoFlex model with a Chamfer-only variant and with variants that removed the global transport matching loss, correspondence anchoring or local motion coherence (Table 4). Ground-truth correspondence labels from the simulations were used only for evaluation and were not supplied to any variant during registration.

**Table 4.** Ablation of the registration constraints in the CryoFlex objective. Ablations were evaluated on the clean TmrAB and PCAT1 simulation-defined benchmarks. Point RMSE measures correspondence-based displacement error at sampled points. Map RMSE measures the distance between the predicted and ground-truth displaced positions of each voxel. Dice@15 measures overlap between predicted and ground-truth masks of the top 15% highest-displacement voxels. The ‘Chamfer-only’ variant uses the same deformation backbone with only the Chamfer distance loss. ‘w/o transport’ removes the global transport matching loss; ‘w/o anchors’ removes correspondence anchoring; ‘w/o LMC’ removes local motion coherence. Best value per dataset and metric is shown in bold. RMSE values are in Å; Dice@15 is dimensionless.

| Dataset | Variant | Point RMSE ↓ | Map RMSE ↓ | Dice@15 ↑ |
| --- | --- | --- | --- | --- |
| TmrAB | Chamfer-only | 0.883 | 0.668 | 0.935 |
|  | w/o transport | 0.798 | 0.597 | 0.938 |
|  | w/o anchors | 0.697 | 0.454 | 0.941 |
|  | w/o LMC | 0.625 | 0.405 | 0.944 |
|  | **Full CryoFlex** | **0.590** | **0.374** | **0.959** |
| PCAT1 | Chamfer-only | 0.976 | 0.627 | 0.932 |
|  | w/o transport | 0.808 | 0.549 | 0.947 |
|  | w/o anchors | 0.676 | 0.433 | 0.955 |
|  | w/o LMC | 0.641 | 0.404 | 0.958 |
|  | **Full CryoFlex** | **0.564** | **0.385** | **0.962** |

Across both datasets, the full CryoFlex model achieved the lowest point RMSE and map RMSE and the highest Dice@15. Among the single-constraint ablations, removing the global transport matching loss produced the largest increase in map RMSE, from 0.374 to 0.597 on TmrAB and from 0.385 to 0.549 on PCAT1. Removing correspondence anchoring produced an intermediate increase in error, whereas removing local motion coherence resulted in a smaller but consistent degradation across the two transitions. The Chamfer-only variant showed the largest errors overall, indicating that the deformation backbone optimized using only the Chamfer distance loss was insufficient to attain the performance of the full model. Because the same kernel regression was applied to all variants, differences in the map-level metrics reflect differences in the displacement vectors at sampled points rather than differences in the projection procedure. The same ordering was observed across the additional nearest-neighbor and map-level metrics reported in Supplementary Table S6. These results show that transport matching, correspondence anchoring and local motion coherence make complementary contributions to CryoFlex performance on these simulation-defined benchmarks.

## 3. Discussion

Reconstructing conformations from particle images and characterizing the differences between selected states are related but distinct steps in cryo-EM analysis. Methods such as cryoDRGN and 3DFlex address the former, whereas CryoFlex addresses the latter (Zhong et al., 2021; Punjani and Fleet, 2023). Given two selected density maps, CryoFlex estimates displacement vectors at sampled points and a voxel-aligned displacement field, whose magnitude defines the flexibility map. This representation makes explicit where corresponding density is displaced, in which direction and by how much, without requiring fitted atomic models to derive the deformation. CryoFlex therefore compares structural motion between selected states directly from cryo-EM density maps.

The density map applications illustrate the analytical value of this density-native representation across different levels of atomic-model availability. In NTCP, the representation localized a gate-opening rearrangement across the transmembrane helices, with paired atomic models available as a consistency reference. Beyond this case, the *α*v*β*8 integrin, tri-snRNP and Spike analyses tested complementary practical settings. In the integrin maps, the same representation characterized motion in a flexible arm for which atomic-model coverage was incomplete. In the tri-snRNP maps, the target reconstruction exhibited attenuated, directionally blurred and poorly defined density in the head region, consistent with the large motion and substantial internal flexibility previously reported for this region (Punjani and Fleet, 2023). After alignment on the comparatively stable body, CryoFlex identified a coherent large-scale head displacement relative to the body. This observation indicates that the displacement representation remained informative where local map quality was degraded, provided that enough density remained shared between the two maps to constrain the correspondence. Applied to successive Spike states selected from an upstream heterogeneous reconstruction, CryoFlex distinguished adjacent intervals of receptor-binding-domain opening and localized weaker coordinated motion elsewhere in the trimer. Together, these cases show that the same representation localized dominant domain rearrangements and secondary regional motions directly in the coordinate system of the source density under incomplete atomic-model coverage and variable local density quality. The accuracy of these estimates was assessed separately under controlled conditions, as discussed below.

The interpretation of the integrin and Spike displacement patterns was further supported by independently reconstructed same-state half-map controls. These controls produced only negligible or low-amplitude, spatially isolated responses, whereas the corresponding conformational-transition pairs retained coherent displacement patterns. Reconstruction-specific variation therefore did not account for the dominant signals in these two cases (Supplementary Fig. S1).

The controlled evaluations first addressed whether CryoFlex estimates the displacement field more accurately than existing registration methods. On the simulation-defined benchmarks, CryoFlex outperformed the strongest registration baseline at both the point and map levels. Because the same kernel regression was applied to every method, the map-level advantage reflected more accurate displacement estimates rather than differences in the voxel-grid projection procedure. Under noisy inputs, absolute displacement errors increased, whereas the spatial localization of high-displacement regions was better preserved, and CryoFlex retained the lowest map-level errors and the highest Dice@15 across the tested SNR range. The registration-constraint ablations showed that global transport matching, correspondence anchoring and local motion coherence made complementary contributions under these benchmark conditions. These results show that CryoFlex provides more accurate displacement recovery than the tested registration methods under both clean and noisy conditions, while regional motion patterns remain informative when map quality limits magnitude accuracy.

The controlled evaluations then addressed whether the estimated displacement corresponds to the between-state displacement of the underlying structures. Against C*α*-referenced endpoint displacements, CryoFlex achieved an endpoint error of approximately 1 Å across nominal resolutions of 3–8 Å (Fig. 6), with a modest system-dependent underestimation of displacement magnitude that did not increase as the nominal resolution became coarser. This agreement shows that, for map pairs simulated independently from paired atomic structures, CryoFlex recovered displacement vectors consistent with the between-state displacement of the underlying atomic structures, rather than only reproducing a simulation-defined voxel displacement field. Together, these controlled evaluations indicate that the CryoFlex displacement representation is quantitatively grounded rather than purely descriptive.

Beyond pairwise motion characterization, the voxel-aligned flexibility map was used to guide downstream structure-processing tasks. Across the integrin, tri-snRNP and Spike applications, the flexibility maps were used to define focused masks for local refinement and to provide spatial priors during cryoDRGN training. CryoFlex-guided local refinement improved the local resolution of the targeted mobile region in all three cases: as a higher gold-standard FSC resolution for the integrin and, at matched gold-standard FSC resolution, as higher estimated local resolution within the mobile regions of tri-snRNP and Spike. Used as a spatial prior during cryoDRGN training, the flexibility map produced better-defined and more continuous density, with fewer fragmented features, than matched baselines.

The present formulation assumes that differences between the two states are predominantly conformational rather than compositional, such that corresponding density features can be related through deformation. CryoFlex does not model the appearance or disappearance of density. Regions arising from compositional differences, from missing structural components or from density that is largely absent in one state therefore cannot be assigned a reliable displacement and remain outside its scope.

Future work could extend CryoFlex from pairwise comparison to the co-ordinated analysis of multiple states, while explicitly distinguishing deformation from density appearance or disappearance. Overall, CryoFlex provides a density-native route from two density maps of distinct states to a quantitative and spatially interpretable description of their pairwise conformational difference. It complements heterogeneous reconstruction by quantifying structural motion between selected states after reconstruction.

## 4. Resource Availability

### 4.1. Lead Contact

Further information and requests for resources should be directed to and will be fulfilled by the Lead Contact, Xiaohua Wan.

### 4.2. Materials Availability

This study did not generate new unique reagents.

### 4.3. Data and Code Availability

The publicly available cryo-EM image datasets analyzed in this study were obtained from the Electron Microscopy Public Image Archive under accessions EMPIAR-10345 (the *α*v*β*8 integrin dataset; https://doi.org/10.6019/EMPIAR-10345) and EMPIAR-10073 (the yeast U4/U6.U5 tri-snRNP dataset; https://doi.org/10.6019/EMPIAR-10073). The CryoBench v1 Spike-MD archive `Spike-MD.zip` (MD5: `6a063a49b1cc7f31e3ad83ced26023b1`) was obtained from Zenodo (https://doi.org/10.5281/zenodo.12528784) (Jeon et al., 2024).

The atomic structures used for the NTCP analyses were obtained from the Protein Data Bank (PDB) under accessions 7PQG (https://doi.org/10.2210/pdb7PQG/pdb) and 7PQQ (https://doi.org/10.2210/pdb7PQQ/pdb) (Goutam et al., 2022). The atomic model used to interpret the *α*v*β*8 integrin analysis was obtained from the PDB under accession 6UJB (https://doi.org/10.2210/pdb6UJB/pdb) (Campbell et al., 2020). The yeast tri-snRNP atomic structure used for anatomical region definition was obtained from the PDB under accession 5GAN (https://doi.org/10.2210/pdb5GAN/pdb) (Nguyen et al., 2016).

The atomic structures used to construct the simulation-defined benchmarks were obtained from the PDB under accessions 6RAH (https://doi.org/10.2210/pdb6RAH/pdb) and 6RAI (https://doi.org/10.2210/pdb6RAI/pdb) for TmrAB (Hofmann et al., 2019), and 7T55 (https://doi.org/10.2210/pdb7T55/pdb) and 7T57 (https://doi.org/10.2210/pdb7T57/pdb) for PCAT1 (Kieuvongngam and Chen, 2022).

The CryoFlex source code and analysis scripts, together with the re-constructed conformational maps, independently reconstructed half-maps, CryoFlex displacement fields and flexibility maps, simulation-defined benchmark datasets, and numerical source data underlying the figures, are available in the project GitHub repository at https://github.com/Dongq1qi/CryoFlex. Any additional information required to reanalyze the data reported in this paper is available from the Lead Contact upon request.

## 5. Acknowledgments

This work was supported in part by the National Natural Science Foundation of China (grant nos. W2511070 and 32241027 to Fa Zhang and 62472034 to Xiaohua Wan, and 62227807 to Bin Hu).

## 6. Author Contributions

Conceptualization, H.D., X.H.W., and F.Z.; Methodology, H.D.; Software, H.D.; Validation, H.D. and X.S.W.; Formal analysis, H.D. and X.S.W.; Investigation, H.D., F.L., Y.C., S.T., C.J., and Z.L.; Data curation, H.D.; Visualization, H.D.; Writing – original draft, H.D.; Writing – review & editing, H.D., F.L., X.H.W., and F.Z.; Supervision, X.H.W., F.Z., and B.H.; Project administration, X.H.W. and F.Z.; Funding acquisition, X.H.W., F.Z., and B.H.

## 8. STAR Methods

### 8.1. Density-native deformation formulation

Let *D*_*s*_ : Ω → ℝ and *D*_*t*_ : Ω → ℝ denote two rigidly aligned cryo-EM density maps of the same macromolecular complex in distinct conformations, defined on a common spatial domain Ω ⊂ ℝ^3^. We formulate their pairwise comparison as a density-native deformation estimation problem: the objective is to estimate a mapping Φ : Ω → Ω that assigns each location *x* in the source map to its corresponding location Φ(*x*) in the target map. Because Φ is not uniquely determined by density similarity alone, we seek an admissible deformation that is consistent with both maps while remaining spatially coherent:

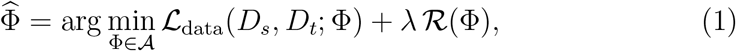

where *A* is the space of admissible non-rigid deformations, ℒ_data_ measures agreement between the source density deformed by Φ and the target density, ℛ enforces spatial coherence on the estimated motion, and *λ >* 0 balances the two terms. The estimated mapping defines the displacement field

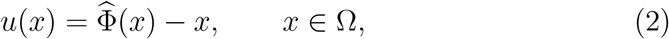

where *u*(*x*) ∈ ℝ^3^ points from *x* to the corresponding location in the target map and therefore carries both the direction and the magnitude of the estimated displacement. Its magnitude

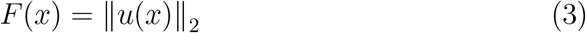

is the local displacement magnitude between the two reconstructions at each location in Ω, and is specific to the pair of states supplied as input. The workflow that implements this formulation is summarized in Fig. 1.

### 8.2. Density-aware point cloud representation

Comparing density values directly on the voxel grid is a brittle way to instantiate the data term ℒ_data_ in Eq. 1, because cryo-EM maps are affected by noise, spatially varying resolution and weak or attenuated density in flexible regions. Motivated by the success of density-vector representations in cryo-EM map alignment (Han et al., 2021; He et al., 2024), we instead represent each density map as a density-aware point cloud: each map contributes a coordinate set 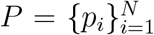 together with attributes derived from its own local density distribution.

For either density map *D* : Ω → ℝ, *P* is sampled inside the molecular contour mask with uniform spacing *s* in physical coordinates. For each sampled point *p*, density values within a local neighborhood *N* (*p*) ⊂ Ω are aggregated with the isotropic Gaussian weight 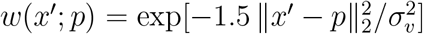, where *σ*_*v*_ sets the spatial scale. The density score and the density-weighted centroid at *p* are

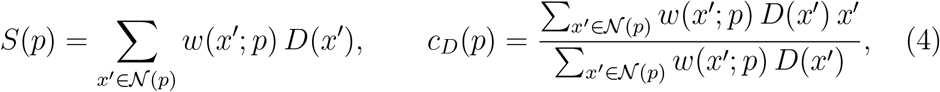

and the unit density vector is

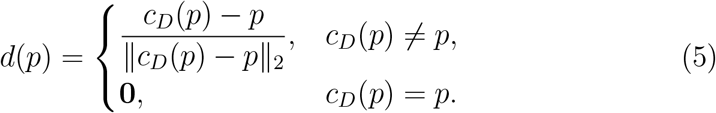

The pair (*S*(*p*), *d*(*p*)) describes the local density and orientation pattern around *p* and is less sensitive to individual voxel noise than raw intensity.

To identify stable structural landmarks, mean shift (Comaniciu and Meer, 2002) was iterated on the sampled points to convergence by repeatedly replacing each point with the density-weighted centroid *c*_*D*_ of its neighborhood. The converged endpoints indicate local density modes and were clustered in three dimensions with DBSCAN (Ester et al., 1996), and the centroid of each cluster was taken as a keypoint. For each keypoint we computed a SHOT descriptor (Tombari et al., 2010) from the neighboring sampled points, which aggregates the spatial distribution and orientation statistics of their density vectors into a rotation-aware histogram. Candidate matches were obtained by nearest-neighbor search in descriptor space and retained only if mutually nearest. Because keypoints are cluster centroids and need not belong to the sampled point sets, each matched keypoint was assigned to its nearest sampled point in the corresponding map, giving a set *C* of matched index pairs between the source and target sampled points that serve as sparse geometric anchors during deformation estimation.

### 8.3. Hierarchical non-rigid deformation

Given the source and target point sets 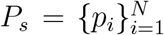 and 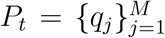, CryoFlex estimates a non-rigid deformation that maps the source sampled points toward the target sampled points. This point-domain estimator instantiates Eq. 1: the matching terms approximate density agreement, while the sparse anchors and local motion coherence (LMC) regularization constrain the otherwise ill-posed deformation.

Inspired by the neural deformation pyramid (Li and Harada, 2022), we parameterize the deformation by a hierarchical network that refines the source coordinates from coarse to fine over *L* levels. With 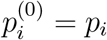, level *l* = 1, …, *L* applies an incremental deformation

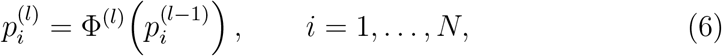

where Φ^(*l*)^ : ℝ^3^ → ℝ^3^ is a neural deformation module. For a point with coordinates (*x, y, z*), level *l* uses the frequency 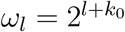 and the six-dimensional positional encoding

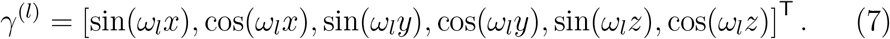

Because the frequency increases with level, coarse levels capture smoother global motion and finer levels resolve more local displacement. The composition Φ^(*L*)^ ◦ · · · ◦ Φ^(1)^ realizes on the sampled points the estimate Φ of Eq. 1.

Each level is optimized with the composite objective

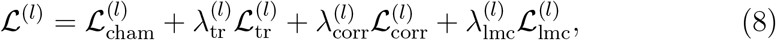

whose weights and level-wise use are reported in the implementation details. The Chamfer distance loss provides a local geometric data term between the deformed source point set 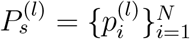 and the target point set:

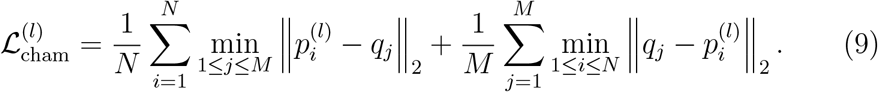

Its nearest-neighbor assignments are made independently and therefore do not impose a globally consistent correspondence between the two point sets. A global transport matching loss addresses this limitation by estimating a soft coupling over all source–target point pairs. We represent 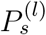 and 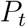 as the empirical measures *α*^(*l*)^ and *β*, each assigning uniform mass to its points, and define the entropically regularized transport cost between them as

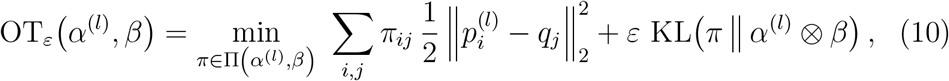

where Π(*α*^(*l*)^, *β*) is the set of couplings whose marginals are *α*^(*l*)^ and *β*, so that *π* transports the full point mass of one measure onto the other, and 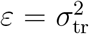 is set by the transport smoothing scale *σ*_tr_ *>* 0. The transport matching loss is the debiased transport divergence

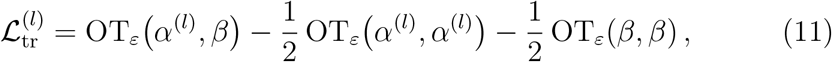

in which the two self-transport terms are obtained from the same expression with both arguments taken from the same point set. Subtracting them removes entropic bias, so identical measures have zero divergence. All three terms are computed in the log domain for numerical stability. This coupling supplies a global distribution-matching signal and is distinct from the sparse anchors below.

To use the sparse anchors, we add the uniformly weighted correspondence loss

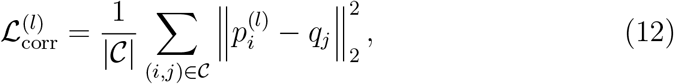

where *C* contains the anchor index pairs (*i, j*) from Sec. 8.2, each linking source point *p*_*i*_ to target point *q*_*j*_. The term uses these pairs as a prior on the deformation: at the coarse levels, where the deformed source is still far from the target and nearest-neighbor geometry alone is ambiguous, they keep the estimate anchored to plausible alignments and stabilize the early stage of optimization. Finally, the LMC term penalizes displacement discrepancies between neighboring sampled points:

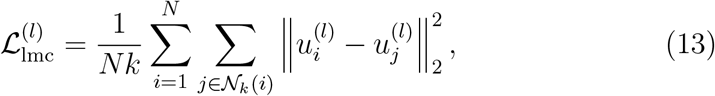

where 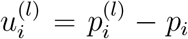 is the accumulated displacement of sampled point *p*_*i*_ at level *l* and *N*_*k*_(*i*) contains its *k* nearest neighbors in the original source point set, with uniform edge weights. It encourages nearby structural regions to move coherently while still allowing spatially varying deformation across domains or flexible segments. After optimization, the deformed position of each sampled point is 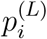, and the corresponding displacement *u*_*i*_ in physical coordinates follows from the coordinate restoration given in the implementation details.

### 8.4. Kernel regression onto the voxel grid

To express the point-domain estimate on the source-map voxel grid, we construct a voxel-aligned displacement vector field before extracting displacement magnitude. For a voxel center *x* ∈ Ω, the displacement vector is estimated by kernel regression over the source sampled points within radius *R* of *x*:

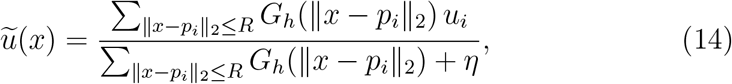

where *G*_*h*_(*r*) = exp[− *r*^2^*/*(2*h*^2^)] is a Gaussian kernel with bandwidth *h* and *η >* 0 prevents division by zero, so that nearby sampled points contribute more strongly and distant points are attenuated. The field 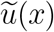 is the kernel-regressed approximation of *u*(*x*) on the voxel grid, and differs from *u*_*i*_ at the sampled points themselves because each value averages the displacements of neighboring points. The reported voxel-level estimate of *F* (*x*) is

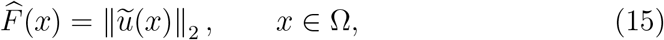

with a molecular mask obtained by thresholding the source density applied so that 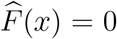 outside the molecular envelope. The reported flexibility map therefore expresses estimated local displacement magnitude between the two states on the voxel grid and remains directly aligned with the source density.

### 8.5. Implementation details

Point clouds were sampled at a default spacing of *s* = 4 Å, with the molecular contour mask of Sec. 8.2 defined separately for each map from its own density distribution. Density-vector construction used *d*_res_ = 16 Å with *σ*_*v*_ = *d*_res_*/*2 = 8 Å, aggregation being truncated to an axis-aligned window extending 2*d*_res_ from each sampled point along each coordinate. Mean shift used an independent bandwidth of 3.0 Å, a 17-voxel window, a step size of 0.05, a convergence threshold of 1.87 × 10^−4^ on the mean squared point update and at most 2,000 iterations. DBSCAN used a distance tolerance equal to *s* and a minimum cluster size of one; SHOT descriptors used a support radius of 7*s*.

Both point clouds were placed on a common numerical scale by a shared source-derived transformation, using the source centroid *c*_*s*_ and radius *r*_*s*_ = max_*i*_ ∥*p*_*i*_ − *c*_*s*_∥_2_:

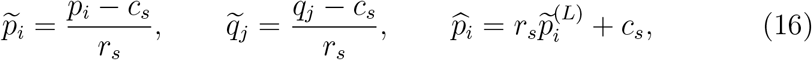

where 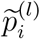 denotes the level-*l* iterate 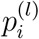 of Sec. 8.3 expressed in these normalized coordinates, 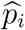 is the recovered physical position of source point *p*_*i*_, and 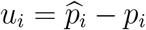 is its displacement in physical coordinates. All deformation and loss evaluations were performed in this normalized system, and physical coordinates were restored only after optimization.

The hierarchical network used *L* = 9 levels and a positional-encoding base exponent of *k*_0_ = −5. Each level contained a three-layer multilayer perceptron of width 128 with ReLU activations, whose outputs parameterized local SE(3) updates as Euler rotations and translations and were scaled by 0.01 so that optimization started near the identity deformation. The deformation modules were optimized independently for each map pair and no pretrained weights were used. Levels were optimized sequentially with Adam at a learning rate of 0.01 for at most 500 iterations each; during optimization of level *l* only that module was trainable, all other levels were frozen, and the coordinates produced at the end of level *l* were detached from its optimization graph and used as the input to level *l* + 1. Optimization stopped when the total loss fell below 10^−4^ or after a relative loss change below 10^−4^ had been observed on 30 optimization checks. Default weights for the Chamfer, transport, anchor and LMC terms were 1, 100, 1 and 1; anchors were active in the first two levels; the LMC graph used 32 nearest neighbors. The transport term used a quadratic ground cost and a smoothing scale of *σ*_tr_ = 0.1 in normalized coordinates, giving *ε* = 0.01, and its weight was set to 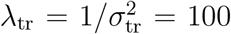 to keep the weighted term numerically comparable to the others. For voxel projection we used *R* = 2.5*s* by default, adjusted within 2*s* ≤ *R* ≤ 3*s* according to point coverage, with *h* = *R/*3.

For the downstream local refinement analyses, a focused mask was constructed automatically from the predicted flexibility map for cryoSPARC Local Refinement. Density support was defined by the 85th percentile of positive density values and expanded by four voxels. Within this support, the flexibility map was Gaussian-smoothed with a standard deviation of 1.5 voxels; voxels above the 95th percentile of the smoothed distribution served as high-confidence seeds and were expanded by connected region growing within the region above the 90th percentile. Seed-containing connected regions were retained, and the dominant component was morphologically cleaned, dilated by eight voxels and padded with a 12-voxel cosine soft edge. Within each dataset, all local refinement conditions used the same particle inputs and default cryoSPARC settings; only the supplied mask differed. These parameters were selected empirically and were not systematically optimized, and should be adapted to the characteristics of each dataset.

For the cryoDRGN analyses, the standard training objective ℒ_cryoDRGN_ (Zhong et al., 2021) was augmented with an auxiliary consistency term between a fixed reference volume *V*_ref_ and the volume *V*_*θ*_ decoded at the latent origin. This term is a voxel-wise weighted mean squared error,

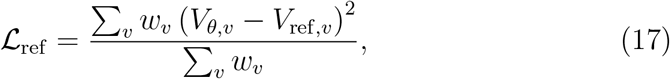

where *v* indexes voxels, *V*_*θ*_ and *V*_ref_ are compared after a global intensity standardization, and each voxel carries its own weight *w*_*v*_ derived from the CryoFlex flexibility map, large where the map predicts little motion and small where it predicts large motion. The two terms were combined as

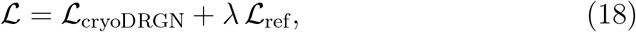

with the global scalar *λ* set to 1 in all runs, so that the spatially varying strength of the constraint is carried entirely by the voxel weights *w*_*v*_: the reconstruction is held close to the reference and retains its frequency content in regions expected to remain static, while conformational variability in the mobile regions is not suppressed. Within each dataset, the baseline and regularized runs used the default network architecture, a latent dimension of 8 and 30 training epochs, with inputs, preprocessing and all remaining training settings held fixed and the CryoFlex-guided regularization as the only difference.

### 8.6. Evaluation protocols and metrics

Two complementary simulation protocols were used. For the C*α*- referenced evaluation, source and target density maps were simulated independently from the pre-aligned paired atomic states of TmrAB (6RAH → 6RAI) (Hofmann et al., 2019) and PCAT1 (7T55 → 7T57) (Kieu-vongngam and Chen, 2022) with `e2pdb2mrc` in EMAN2 (Tang et al., 2007). Maps were generated at a voxel size of 1.5 Å and nominal resolutions of 3, 4, 5, 6 and 8 Å. Because each map was generated directly from its corresponding atomic state, matched C*α* atoms provided the reference endpoint displacement vectors.

For the simulation-defined voxel-field benchmarks, the source density map was simulated from the pre-aligned source-state atomic structure at a voxel size of 1.5 Å and a nominal resolution of 3 Å. The evaluation target was then generated by warping this source map along a known dense displacement field representing the corresponding between-state motion on the source voxel grid. Because this field was used directly to generate the target map, it is by construction the exact displacement field relating the source and target maps rather than an estimate of an unknown transformation. Its values provided the reference displacement vectors at the source sampled points and the ground-truth displacement field on the voxel grid. This construction shares no kernels, bandwidths or smoothness assumptions with the kernel regression that CryoFlex uses to project displacement vectors onto the voxel grid (Sec. 8.4).

For the noise-robustness experiments, noise was added to each clean benchmark map before point sampling using a density-dependent Gaussian model:

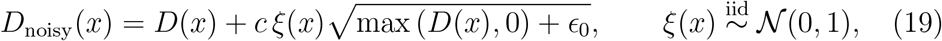

where *ϵ*_0_ is a small numerical constant. The global scale *c* was selected so that the ratio of the summed squared clean density to the summed squared added noise, evaluated over the molecular region of the map, matched the prescribed map-level SNR. We evaluated linear SNR values of 5, 2 and 1.

Every method received the same rigidly aligned source and target point sets sampled from the two density maps, using publicly available baseline implementations with the parameters given in their original reports. As part of CryoFlex, density scores and unit density vectors were computed from the same maps to construct density-informed correspondence anchors, whose contribution was evaluated in the ablation analysis (Table 4).

All methods were evaluated against the same clean targets, with their point displacements projected to the voxel grid by the same kernel regression. The reference displacement vectors at the sampled points, the known source– target correspondences and the ground-truth voxel-grid field were never provided to any method.

Point-level metrics evaluate displacement-vector errors at the sampled points: correspondence-based RMSE and MAE, computed from the distance between each predicted point position and its known ground-truth target position; one-sided nearest-neighbor RMSE and MAE (RMSENN and MAENN), computed from the distance from each predicted point to the nearest point of the clean target set without using correspondence labels; the symmetric Chamfer-*L*_1_ distance, taken as the sum of the two directional mean nearest-neighbor distances; and HD95, taken as the maximum of the two directional 95th-percentile Hausdorff distances.

Voxel-level metrics were computed within the molecular region of the reference map. Map RMSE, MAE and RMSE@15 evaluate displacement-vector errors, computed from the distance between the predicted and ground-truth displaced positions of each voxel; Corr, Dice@15 and AUPRC@15 evaluate displacement-magnitude agreement and the localization of high-displacement regions. Corr is the Pearson correlation between the predicted and ground-truth displacement magnitudes. The ground-truth and predicted high-displacement regions were each obtained by rank-thresholding their own displacement magnitudes at the top 15% and retaining all tied voxels; Dice@15 measures the overlap of these two regions, RMSE@15 is restricted to the ground-truth region, and AUPRC@15 scores the ranking of ground-truth high-displacement voxels by predicted magnitude with that region as the positive class. All distance-based metrics are reported in Å.

For paired states with matched C*α* atoms, the between-state reference displacement vector of each atom was taken as the difference between its target- and source-state coordinates, and the corresponding CryoFlex estimate was obtained by evaluating the kernel-regressed field *ũ* of Eq. 14 at the source-state C*α* position. Agreement was quantified by the endpoint error (EPE) and RMSE between the two vector fields, both of which include the directional component, and by the Pearson correlation and mean signed bias, predicted minus reference, of their magnitudes. Cross-resolution consistency was calculated as the Pearson correlation between displacement magnitudes predicted at the same C*α* positions from the 3- and 8-Å maps. EPE, RMSE and signed bias are reported in Å; Pearson correlation is dimensionless.

For the same-state half-map controls, voxel-level statistics were computed over the common molecular region of the two half-maps. For each displacement threshold *τ* ∈ {1, 2, 4, 8} Å, the active voxel set and false-active rate were

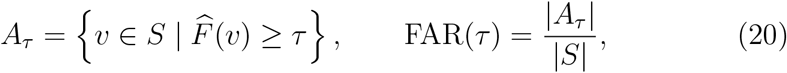

where |*S*| denotes the number of voxels in the common molecular region. To quantify the spatial extent of the suprathreshold response, *A*_*τ*_ was partitioned into three-dimensional connected components using 26-neighbor connectivity. With *C*_max_(*τ*) the number of voxels in the largest component, the largest-component fraction of the region was

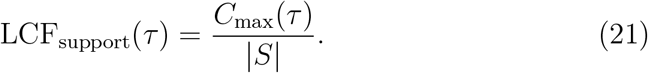

Comparisons of spatial extent were reported at *τ* = 4 Å.

Gold-standard Fourier shell correlation (FSC) was calculated in cryoSPARC between the two independently reconstructed half-maps. Reported resolutions are the reciprocal of the spatial frequency at which the mask-corrected gold-standard FSC curve first crosses the 0.143 threshold.

## Supplementary Information

Independent half-map controls distinguish reconstruction variability from conformational change in two datasets

Because CryoFlex estimates a displacement field for any pair of input maps, the existence of an estimated displacement field alone cannot distinguish conformational change from density differences introduced by independent reconstruction. To determine whether such reconstruction variability could be misinterpreted as conformational change, we constructed same-state controls using independently reconstructed half-maps generated in cryoSPARC for EMPIAR-10345 and Spike. The two half-maps in each control were reconstructed from independent particle subsets but represented the same underlying conformational ensemble. Displacements estimated between them therefore provided a direct measure of the background motion signal arising from reconstruction variability. CryoFlex was applied to each half-map pair using the same analysis pipeline as for the corresponding conformational-transition pairs. We quantified the resulting background signal using the false-active rate (FAR), defined as the fraction of voxels within the common density support whose estimated displacement exceeded 1, 2, 4 or 8 Å.

CryoFlex clearly separated same-state half-map variation from genuine conformational change in both datasets (Fig. S1). For EMPIAR-10345, the FAR of the half-map control decreased rapidly from 0.306% at 1 Å to 0.070% at 2 Å and 0.0026% at 4 Å, with no voxels exceeding 8 Å. By comparison, the corresponding conformational-transition pair retained FAR values of 6.295%, 5.388%, 4.537% and 3.521% at the same thresholds, respectively. The spatial organization of the detected signal provided an even clearer distinction. At the 4 Å threshold, the largest connected component in the half-map control occupied only 0.0026% of the common density support, compared with 4.515% for the conformational-transition pair. Thus, the residual response between the EMPIAR-10345 half-maps consisted of sparse, low-amplitude fluctuations rather than the spatially coherent deformation pattern observed for conformational change.

The separation was even more pronounced for Spike. No active voxels were detected in the half-map control at any of the four displacement thresholds, whereas the conformational-transition pair produced FAR values of 12.056%, 9.402%, 6.060% and 2.532% at 1, 2, 4 and 8 Å, respectively. Its largest connected component at 4 Å occupied 3.004% of the common density support, in contrast to the absence of any suprathreshold component in the half-map control. The lower background response in the Spike dataset is consistent with its high-resolution reconstruction (3.06 Å), for which the independently reconstructed half-maps retain more closely matched structural density and therefore contain less reconstruction-specific variation that could be interpreted as motion. Together, the magnitude and spatial-connectivity analyses distinguish isolated fluctuations arising from independent reconstruction from spatially coherent deformation associated with conformational change in these two datasets.

Local motion coherence regularization (LMC) further suppressed isolated false-positive displacements in the more challenging EMPIAR-10345 control (Fig. S1d). Removing LMC increased the FAR at 4 Å from 0.010% to 0.156%, corresponding to an approximately 15-fold increase, and introduced voxels with estimated displacements exceeding 8 Å. The maximum estimated displacement simultaneously increased from 4.38 Å to 14.79 Å. The same full pipeline containing LMC retained the strong, spatially extended signals observed for the conformational-transition pairs. The low FAR of the half-map controls therefore did not arise from uniformly suppressing the estimated displacement signal. Instead, these results indicate that LMC limited the amplification of local reconstruction fluctuations into high-magnitude, spatially incoherent displacement estimates while preserving transition-level deformation.

**Figure S1.**
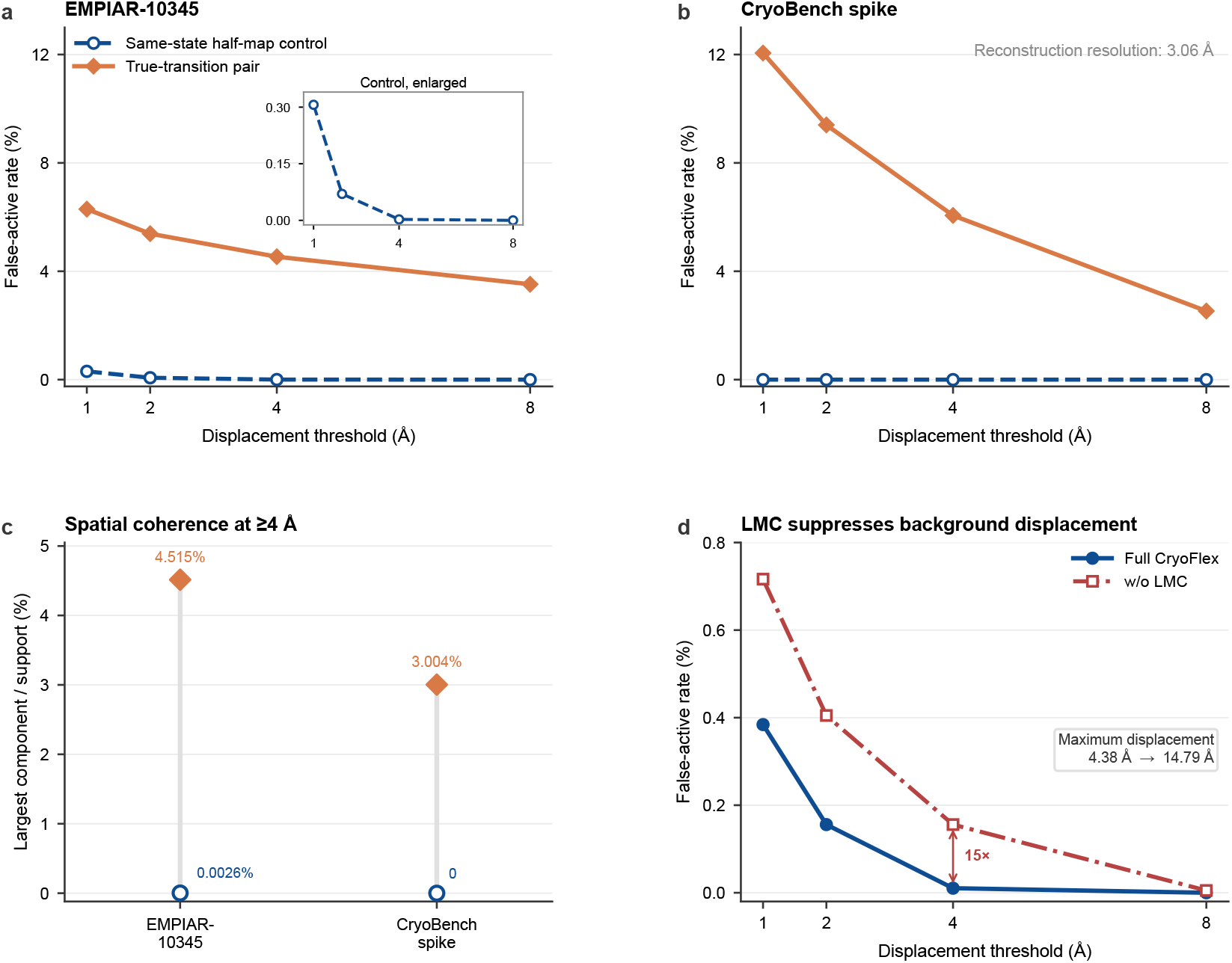
Independent half-map controls distinguish reconstruction variability from conformational change in two datasets. **a,b**, False-active rate (FAR), defined as the fraction of voxels within the common density support whose estimated displacement exceeded the indicated threshold, for same-state half-map controls and corresponding conformational-transition pairs from EMPIAR-10345 (**a**) and Spike (**b**). The inset in **a** enlarges the low-FAR range of the EMPIAR-10345 half-map control. **c**, Fraction of the common density support occupied by the largest connected component of voxels with estimated displacement ≥ 4 Å. The half-map controls produced either no suprathreshold component or only a negligible spatially isolated component, whereas the conformational-transition pairs retained spatially extended deformation patterns. **d**, Effect of local motion coherence regularization (LMC) on the EMPIAR-10345 half-map control. Removing LMC increased the FAR across displacement thresholds, produced an approximately 15-fold increase at 4 Å and increased the maximum estimated displacement from 4.38 Å to 14.79 Å. The strong signals retained by the full CryoFlex pipeline for the conformational-transition pairs in **a,b** show that suppression of the half-map background did not arise from uniformly eliminating transition-level deformation.

### Gold-standard FSC assessment of tri-snRNP local refinements

To complement the spatial local-resolution comparison in Fig. 4c, we evaluated the gold-standard FSC resolution of the three tri-snRNP local refinements using the same procedure. The original, CryoFlex-guided and global-mask local refinement conditions each reached a resolution of 4.24 Å after FSC mask correction (Fig. S2). Thus, the three conditions reached comparable gold-standard FSC resolutions, while the FSC-based local-resolution maps in Fig. 4c distinguished their spatial resolution distributions within the mobile region.

**Figure S2.**
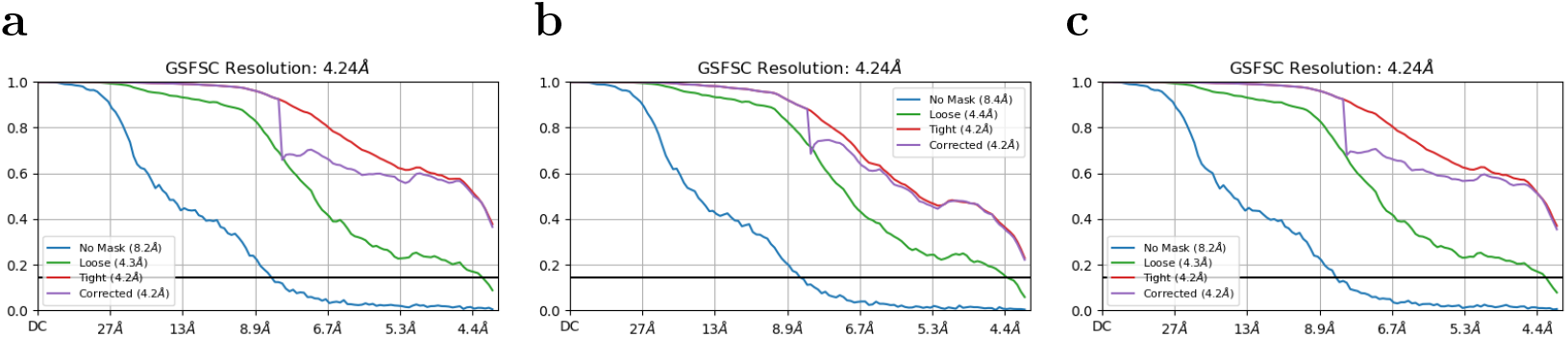
Gold-standard FSC assessment of tri-snRNP local refinements. **a–c**, Gold-standard FSC curves for the original (**a**), CryoFlex-guided (**b**) and global-mask (**c**) local refinement conditions. Curves show the unmasked, loose-mask, tight-mask and mask-corrected correlations generated during FSC assessment. All three refinement conditions reached a final gold-standard FSC resolution of 4.24 Å.

### Gold-standard FSC assessment of Spike local refinements

To complement the spatial local-resolution comparison in Fig. 5d, we evaluated the gold-standard FSC resolution of the three Spike local refinements using the same procedure. The original, CryoFlex-guided and global-mask local refinement conditions each reached a resolution of 3.06 Å after FSC mask correction (Fig. S3). Thus, the three conditions reached comparable gold-standard FSC resolutions, while the FSC-based local-resolution maps in Fig. 5d distinguished their spatial resolution distributions within the mobile region.

**Figure S3.**
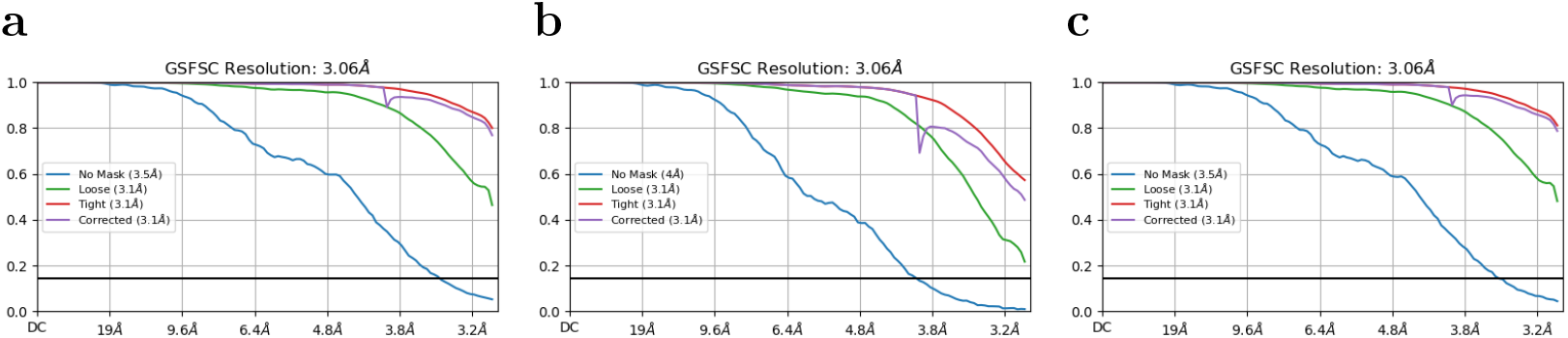
Gold-standard FSC assessment of Spike local refinements. **a–c**, Gold-standard FSC curves for the original (**a**), CryoFlex-guided (**b**) and global-mask (**c**) local refinement conditions. Curves show the unmasked, loose-mask, tight-mask and mask-corrected correlations generated during FSC assessment. All three refinement conditions reached a final gold-standard FSC resolution of 3.06 Å.

### Full registration comparison on the clean benchmarks

Table S1 extends the main registration comparison (Table 1) to all methods and to additional metrics that are standard in the point cloud registration literature. RMSE and MAE are correspondence-based; RMSENN and MAENN are their nearest-neighbor counterparts; CD is the symmetric Chamfer-*L*_1_ distance defined as the sum of the two directional mean nearest-neighbor distances, and HD95 is the maximum of the directional 95th-percentile nearest-neighbor distances. All distance values are in Å, and lower is better. NSFP is included here; its large-scale point cloud inversion produced very large correspondence-based RMSE and MAE despite moderate nearest-neighbor errors. We therefore report it as a diagnostic case rather than include it in the main comparative conclusions.

### Full voxel-grid displacement comparison on the clean benchmarks

Table S2 extends the main voxel-grid displacement comparison (Table 2) to all methods and adds MAE and AUPRC@15. RMSE, MAE and RMSE@15 are in Å; Corr, Dice@15 and AUPRC@15 are dimensionless. Corr is the voxelwise Pearson correlation between the predicted flexibility map and the ground-truth displacement-magnitude map. Negative values indicate an inverse spatial association between the predicted and ground-truth displacement patterns rather than an invalid measurement. AUPRC@15 is the area under the precision–recall curve for the top 15% highest-displacement voxels.

**Table S1.**
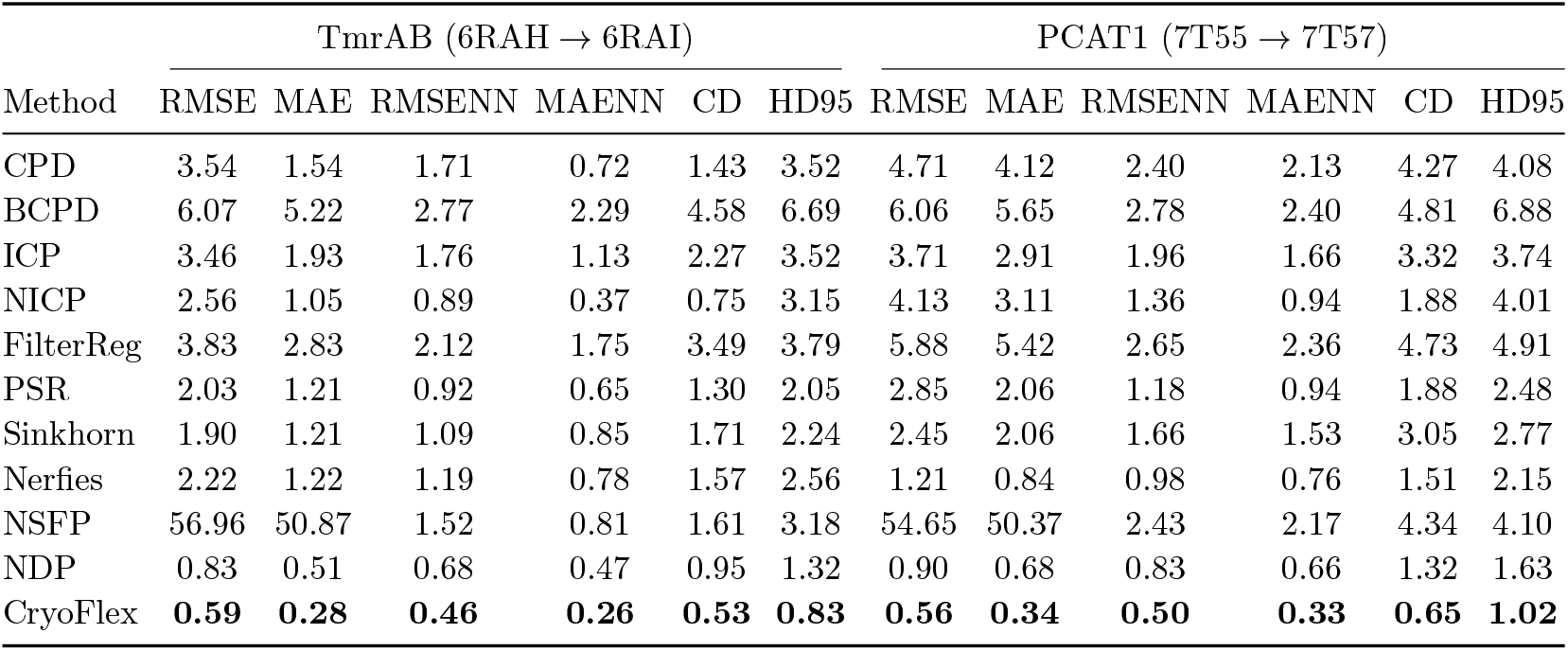
Registration accuracy on the clean benchmarks (all methods). RMSE/MAE are correspondence-based, RMSENN/MAENN are nearest-neighbor based, CD is the symmetric Chamfer-*L*_1_ distance and HD95 is the 95th-percentile Hausdorff distance, as defined in STAR Methods. All distance values in Å; lower is better. Best value per column in bold.

### Full noise-robustness comparison

Tables S3–S5 report the voxel-grid displacement accuracy of all methods under noisy density-derived inputs at signal-to-noise ratios of 5, 2 and 1, extending the main-text comparison (Table 3), which lists only CryoFlex and the two closest baselines. Columns match the clean-input table (Table S2). CryoFlex achieved the best value of every metric on both benchmarks at all three noise levels. Under all noisy-input conditions, CPD produced a spatially constant zero-displacement map. Pearson Corr was therefore undefined because the predicted displacement magnitudes had zero variance and is reported as −−.

### Additional metrics for the registration-constraint ablation

Table S6 extends the main registration-constraint ablation (Table 4) with nearest-neighbor error between point sets, map correlation, high-displacement-region error and threshold-free high-displacement localization. These metrics were not included in the main table to keep the constraint comparison focused on point RMSE, map RMSE and Dice@15.

**Table S2.** Flexibility-map accuracy on the clean benchmarks (all methods). RMSE, MAE and RMSE@15 are in Å; Corr, Dice@15 and AUPRC@15 are dimensionless. Best value per column in bold.

| Method | TmrAB (6RAH → 6RAI) | | | | | | PCAT1 (7T55 → 7T57) | | | | | |
| --- | --- | --- | --- | --- | --- | --- | --- | --- | --- | --- | --- | --- |
|  | RMSE | MAE | Corr | Dice@15 | RMSE@15 | AUPRC@15 | RMSE | MAE | Corr | Dice@15 | RMSE@15 | AUPRC@15 |
| CPD | 3.45 | 1.54 | -0.618 | 0.000 | 8.63 | 0.081 | 4.70 | 4.10 | -0.400 | 0.041 | 8.88 | 0.097 |
| BCPD | 3.32 | 2.95 | 0.708 | 0.695 | 2.95 | 0.783 | 3.15 | 2.69 | 0.408 | 0.240 | 2.00 | 0.220 |
| ICP | 3.07 | 1.46 | 0.444 | 0.461 | 7.68 | 0.370 | 2.28 | 1.45 | 0.547 | 0.388 | 5.26 | 0.317 |
| NICP | 1.94 | 0.81 | 0.852 | 0.672 | 4.89 | 0.782 | 3.04 | 2.20 | 0.363 | 0.369 | 6.30 | 0.418 |
| FilterReg | 2.94 | 1.64 | 0.151 | 0.292 | 7.16 | 0.325 | 3.37 | 2.87 | 0.381 | 0.313 | 2.68 | 0.284 |
| PSR | 0.92 | 0.52 | 0.953 | 0.886 | 1.52 | 0.955 | 1.42 | 0.86 | 0.835 | 0.782 | 3.05 | 0.815 |
| Sinkhorn | 0.87 | 0.49 | 0.972 | 0.905 | 1.96 | 0.970 | 1.37 | 0.99 | 0.876 | 0.893 | 2.54 | 0.898 |
| Nerfies | 1.67 | 0.82 | 0.930 | 0.766 | 4.08 | 0.880 | 0.91 | 0.54 | 0.930 | 0.901 | 1.65 | 0.960 |
| NSFP | 49.99 | 48.83 | 0.515 | 0.480 | 56.86 | 0.452 | 47.55 | 46.72 | 0.253 | 0.135 | 44.45 | 0.168 |
| NDP | 0.51 | 0.27 | 0.986 | 0.946 | 0.87 | 0.986 | 0.52 | 0.34 | 0.975 | 0.960 | 0.77 | 0.983 |
| CryoFlex | **0.37** | **0.19** | **0.992** | **0.959** | **0.74** | **0.992** | **0.38** | **0.23** | **0.986** | **0.962** | **0.52** | **0.991** |

**Table S3.** Flexibility-map accuracy at SNR 5 (all methods). Columns and units as in Table S2. Best value per column in bold.

| Method | TmrAB (6RAH → 6RAI) | | | | | | PCAT1 (7T55 → 7T57) | | | | | |
| --- | --- | --- | --- | --- | --- | --- | --- | --- | --- | --- | --- | --- |
|  | RMSE | MAE | Corr | Dice@15 | RMSE@15 | AUPRC@15 | RMSE | MAE | Corr | Dice@15 | RMSE@15 | AUPRC@15 |
| CPD | 3.63 | 2.17 | -- | 0.261 | 8.77 | 0.150 | 4.76 | 4.17 | -- | 0.261 | 8.94 | 0.150 |
| BCPD | 3.12 | 2.79 | 0.750 | 0.741 | 2.97 | 0.865 | 3.56 | 3.01 | 0.319 | 0.163 | 2.00 | 0.190 |
| ICP | 3.03 | 1.51 | 0.746 | 0.747 | 7.56 | 0.874 | 3.47 | 2.86 | 0.705 | 0.576 | 7.06 | 0.640 |
| NICP | 2.04 | 1.04 | 0.842 | 0.746 | 4.99 | 0.843 | 3.14 | 2.50 | 0.564 | 0.460 | 6.28 | 0.513 |
| FilterReg | 3.14 | 2.32 | -0.320 | 0.002 | 6.85 | 0.092 | 2.67 | 2.23 | 0.330 | 0.233 | 3.60 | 0.216 |
| PSR | 1.18 | 0.81 | 0.918 | 0.832 | 1.62 | 0.895 | 1.60 | 1.15 | 0.755 | 0.701 | 3.12 | 0.762 |
| Sinkhorn | 0.93 | 0.56 | 0.959 | 0.896 | 1.99 | 0.960 | 1.38 | 1.00 | 0.871 | 0.900 | 2.53 | 0.929 |
| Nerfies | 1.66 | 0.96 | 0.917 | 0.757 | 3.89 | 0.867 | 1.38 | 0.94 | 0.872 | 0.861 | 2.36 | 0.929 |
| NSFP | 50.02 | 48.91 | 0.501 | 0.457 | 55.60 | 0.405 | 47.68 | 46.89 | 0.346 | 0.256 | 46.96 | 0.223 |
| NDP | 1.98 | 1.44 | 0.870 | 0.811 | 3.97 | 0.871 | 4.75 | 4.17 | 0.741 | 0.886 | 8.93 | 0.948 |
| CryoFlex | **0.81** | **0.54** | **0.959** | **0.907** | **1.55** | **0.969** | **0.82** | **0.63** | **0.944** | **0.942** | **0.95** | **0.983** |

**Table S4.** Flexibility-map accuracy at SNR 2 (all methods). Columns and units as in Table S2. Best value per column in bold.

| Method | TmrAB (6RAH → 6RAI) | | | | | | PCAT1 (7T55 → 7T57) | | | | | |
| --- | --- | --- | --- | --- | --- | --- | --- | --- | --- | --- | --- | --- |
|  | RMSE | MAE | Corr | Dice@15 | RMSE@15 | AUPRC@15 | RMSE | MAE | Corr | Dice@15 | RMSE@15 | AUPRC@15 |
| CPD | 3.63 | 2.17 | -- | 0.261 | 8.77 | 0.150 | 4.76 | 4.17 | -- | 0.261 | 8.94 | 0.150 |
| BCPD | 6.51 | 5.99 | 0.761 | 0.768 | 7.27 | 0.886 | 3.92 | 3.29 | 0.283 | 0.126 | 1.83 | 0.183 |
| ICP | 3.07 | 1.53 | 0.747 | 0.743 | 7.64 | 0.871 | 3.72 | 3.11 | 0.674 | 0.524 | 7.46 | 0.553 |
| NICP | 2.08 | 1.07 | 0.835 | 0.716 | 5.09 | 0.815 | 3.21 | 2.60 | 0.564 | 0.468 | 6.39 | 0.521 |
| FilterReg | 3.09 | 2.30 | -0.169 | 0.005 | 6.66 | 0.105 | 2.51 | 2.05 | 0.332 | 0.234 | 3.84 | 0.215 |
| PSR | 1.29 | 0.88 | 0.907 | 0.801 | 1.53 | 0.874 | 1.62 | 1.19 | 0.752 | 0.671 | 3.08 | 0.744 |
| Sinkhorn | 0.94 | 0.59 | 0.959 | 0.892 | 1.99 | 0.962 | 1.41 | 1.02 | 0.854 | 0.889 | 2.58 | 0.918 |
| Nerfies | 1.74 | 1.07 | 0.918 | 0.763 | 3.99 | 0.883 | 1.60 | 1.30 | 0.911 | 0.889 | 1.79 | 0.957 |
| NSFP | 49.80 | 48.58 | 0.518 | 0.520 | 57.07 | 0.455 | 47.84 | 46.96 | 0.264 | 0.169 | 45.40 | 0.181 |
| NDP | 2.14 | 1.53 | 0.840 | 0.853 | 4.49 | 0.915 | 4.75 | 4.17 | 0.557 | 0.491 | 8.93 | 0.474 |
| CryoFlex | **0.83** | **0.56** | **0.966** | **0.895** | **1.52** | **0.968** | **0.77** | **0.59** | **0.944** | **0.943** | **1.00** | **0.983** |

**Table S5.** Flexibility-map accuracy at SNR 1 (all methods). Columns and units as in Table S2. Best value per column in bold. At this noise level, NDP produced an inversely associated displacement pattern on PCAT1, with negative Corr and near-zero Dice@15.

| Method | TmrAB (6RAH → 6RAI) | | | | | | PCAT1 (7T55 → 7T57) | | | | | |
| --- | --- | --- | --- | --- | --- | --- | --- | --- | --- | --- | --- | --- |
|  | RMSE | MAE | Corr | Dice@15 | RMSE@15 | AUPRC@15 | RMSE | MAE | Corr | Dice@15 | RMSE@15 | AUPRC@15 |
| CPD | 3.63 | 2.17 | -- | 0.261 | 8.77 | 0.150 | 4.76 | 4.17 | -- | 0.261 | 8.94 | 0.150 |
| BCPD | 7.05 | 6.50 | 0.758 | 0.758 | 8.06 | 0.880 | 4.48 | 3.72 | 0.266 | 0.111 | 1.54 | 0.181 |
| ICP | 3.08 | 1.54 | 0.737 | 0.733 | 7.66 | 0.859 | 3.89 | 3.28 | 0.637 | 0.468 | 7.74 | 0.457 |
| NICP | 2.16 | 1.09 | 0.810 | 0.699 | 5.29 | 0.792 | 3.26 | 2.65 | 0.564 | 0.473 | 6.41 | 0.526 |
| FilterReg | 3.11 | 2.37 | -0.223 | 0.048 | 6.57 | 0.103 | 2.29 | 1.83 | 0.414 | 0.325 | 3.67 | 0.298 |
| PSR | 1.28 | 0.89 | 0.906 | 0.812 | 1.72 | 0.889 | 1.67 | 1.25 | 0.754 | 0.710 | 2.98 | 0.743 |
| Sinkhorn | 0.97 | 0.64 | 0.956 | 0.863 | 1.99 | 0.933 | 1.41 | 1.04 | 0.847 | 0.891 | 2.50 | 0.903 |
| Nerfies | 1.79 | 1.08 | 0.920 | 0.798 | 4.17 | 0.902 | 2.02 | 1.77 | 0.915 | 0.902 | 1.88 | 0.967 |
| NSFP | 49.64 | 48.42 | 0.504 | 0.494 | 56.65 | 0.442 | 48.16 | 47.32 | 0.329 | 0.232 | 47.11 | 0.212 |
| NDP | 3.62 | 2.16 | 0.848 | 0.580 | 8.77 | 0.675 | 4.76 | 4.17 | -0.070 | 0.007 | 8.94 | 0.111 |
| CryoFlex | **0.87** | **0.62** | **0.959** | **0.870** | **1.57** | **0.956** | **0.89** | **0.70** | **0.932** | **0.937** | **0.99** | **0.978** |

**Table S6.** Additional ablation metrics on the clean simulation-defined benchmarks. RMSENN is the nearest-neighbor geometric error between the deformed source point set and the target point set. Corr denotes the Pearson correlation between the predicted flexibility map and the ground-truth displacement-magnitude map. RMSE@15 is the map error computed within the ground-truth top 15% highest-displacement mask. AUPRC@15 is the area under the precision-recall curve for identifying the same high-displacement mask. RMSENN and RMSE@15 are in Å; Corr and AUPRC@15 are dimensionless. Best value per dataset and metric is shown in bold.

| Dataset | Variant | RMSENN | Corr | RMSE@15 | AUPRC@15 |
| --- | --- | --- | --- | --- | --- |
| TmrAB | Chamfer-only | 0.768 | 0.972 | 0.989 | 0.971 |
|  | w/o transport | 0.680 | 0.974 | 0.894 | 0.975 |
|  | w/o anchors | 0.609 | 0.989 | 0.884 | 0.987 |
|  | w/o LMC | 0.550 | 0.991 | 0.776 | 0.989 |
|  | Full CryoFlex | **0.461** | **0.992** | **0.742** | **0.992** |
| PCAT1 | Chamfer-only | 0.871 | 0.967 | 0.718 | 0.975 |
|  | w/o transport | 0.777 | 0.980 | 0.684 | 0.984 |
|  | w/o anchors | 0.644 | 0.982 | 0.620 | 0.990 |
|  | w/o LMC | 0.610 | 0.985 | 0.572 | 0.991 |
|  | Full CryoFlex | **0.504** | **0.986** | **0.516** | **0.992** |

